# Allosteric Competition and Inhibition in AMPA Receptors

**DOI:** 10.1101/2023.11.28.569057

**Authors:** W. Dylan Hale, Alejandra Montaño Romero, Cuauhtemoc U. Gonzalez, Vasanthi Jayaraman, Albert Y. Lau, Richard L. Huganir, Edward C. Twomey

## Abstract

Excitatory neurotransmission is principally mediated by AMPA-subtype ionotropic glutamate receptors (AMPARs). Dysregulation of AMPARs is the cause of many neurological disorders and how therapeutic candidates such as negative allosteric modulators inhibit AMPARs is unclear. Here, we show that non-competitive inhibition desensitizes AMPARs to activation and prevents positive allosteric modulation. We dissected the noncompetitive inhibition mechanism of action by capturing AMPARs bound to glutamate and the prototypical negative allosteric modulator, GYKI-52466, with cryo-electron microscopy. Noncompetitive inhibition by GYKI-52466, which binds in the transmembrane collar region surrounding the ion channel, negatively modulates AMPARs by decoupling glutamate binding in the ligand binding domain from the ion channel. Furthermore, during allosteric competition between negative and positive modulators, negative allosteric modulation by GKYI-52466 outcompetes positive allosteric modulators to control AMPAR function. Our data provide a new framework for understanding allostery of AMPARs and foundations for rational design of therapeutics targeting AMPARs in neurological diseases.

## Introduction

The human brain consists of roughly 100 billion neurons^1^. Central to the brain’s function is the ability of neurons to communicate with one another^2^. Most of the communication between neurons occurs at contact sites called synapses. There are over 100 trillion synaptic connections between neurons in the brain, and most of these synapses are glutamatergic, where the neurotransmitter glutamate (Glu) is released by a pre-synaptic neuron and received by a post-synaptic neuron^3^. Ionotropic glutamate receptors (iGluRs) in the membrane of the post-synaptic neuron bind Glu and allow positively charged ions to enter the post-synaptic neuron, which depolarizes the post-synaptic membrane^4^. This excites a signaling cascade in the post-synaptic neuron that enables signaling to be transduced to other neurons and is vital for information processing. Specialized iGluRs, α-amino-3-hydroxy-5-methyl-4-isoxazolepropionic acid receptors (AMPARs), initiate the depolarization of the post-synaptic neuron and contribute to the activation of the other iGluR subtypes^2^. AMPARs mediate the majority of fast, synchronous excitation in the vertebrate brain and proper signaling of AMPARs in the post-synaptic membrane is required for healthy brain function.

Dysregulation of AMPARs is a major contributor to many neurological disorders. These include schizophrenia, anxiety, chronic pain, epilepsy, learning impairment, Alzheimer’s, and Parkinson’s^4^. A principal approach to therapeutically targeting AMPARs is through allosteric modulation with small molecules. Allosteric modulators enable AMPAR function to be positively or negatively tuned from binding sites independent of Glu binding. Such molecules are clinically promising with broad applications across neurological disorders. However, despite the central role of AMPARs in synaptic signaling and their roles in human diseases, only a single molecule, perampanel (Fycompa®), is approved by the US FDA for targeting AMPARs for therapeutic benefit^4,5^. Perampanel is approved by the FDA to treat epilepsy^6^, and perampanel/perampanel-like molecules (PPLMs) also show promise in treating chronic pain. PPLMs belong to a class of noncompetitive inhibitors typified by the prototype compound 4-(8-Methyl-9*H*-1,3-dioxolo[4,5- *h*][2,3]benzodiazepin-5-yl)-benzenamine dihydrochloride (GYKI-52466)^4,7,8^, which binds to the AMPAR transmembrane domain (TMD)^9^. PPLMs exert characteristically similar effects on AMPAR function and bind to the same site in the TMD^5,8–11^. In the presence of PPLMs, AMPARs show a marked reduction in ion channel conductance irrespective of agonist concentration, and PPLMs inhibit channel function irrespective of channel state or membrane voltage^5,8–11^. PPLMs are effective at reducing epileptic behavior in mice and *in vitro*^12,13^, but perampanel treatment in humans produces undesirable side effects such as dizziness, somnolence, and ataxia^14^, underscoring the need for refined AMPAR inhibitors for treating neurological disorders. While the binding sites of PPLMs have been generally described^9^, the mechanism by which PPLMs inhibit AMPAR function is unknown. This is a major roadblock in therapeutically targeting AMPARs with improved inhibitors.

AMPARs are tetrameric ligand-gated ion channels, made up of GluA1-4 subunits, encoded by the *Gria1-4* genes^4^. AMPARs couple extracellular binding of Glu to ion flux across the postsynaptic membrane by physically linking four extracellular clamshell-shaped ligand binding domains (LBDs) to transmembrane (TM) helices that occlude entry to the cation channel (**Fig. 1a**)^15,16^. Glu binding to the AMPAR LBDs initiates the gating cycle in which the receptors transition through their main functional states: resting, activated, and desensitized^4,17^. Activation begins with Glu binding, which drives the lower half of the LBD clamshell (D2) closer to the upper half (D1) of the LBD, closing the LBD clamshell-like structure around the neurotransmitter (**Fig. 1a**)^15,16^. AMPAR LBDs locally dimerize within the tetrameric receptor, and coordinated clamshell closure maximizes the interface between the upper D1 lobes of LBD dimer pairs (**Fig. 1a**). Movement of the D2 LBD lobes following Glu binding activates the receptor by pulling open the M3 TM helices that define the receptor pore along with the M2 helices, which define the selectivity filter, enabling cation influx. However, the active state is short-lived, and the receptor quickly desensitizes following activation to protect the neuron from toxic influx of cations. Desensitization occurs when the LBDs roll away from each other, which ruptures the D1 interface and closes the ion channel below and reduces separation between the LBD D2’s. AMPARs return to their resting state after unbinding Glu (**Fig. 1a**)^18^.

**Figure 1.**
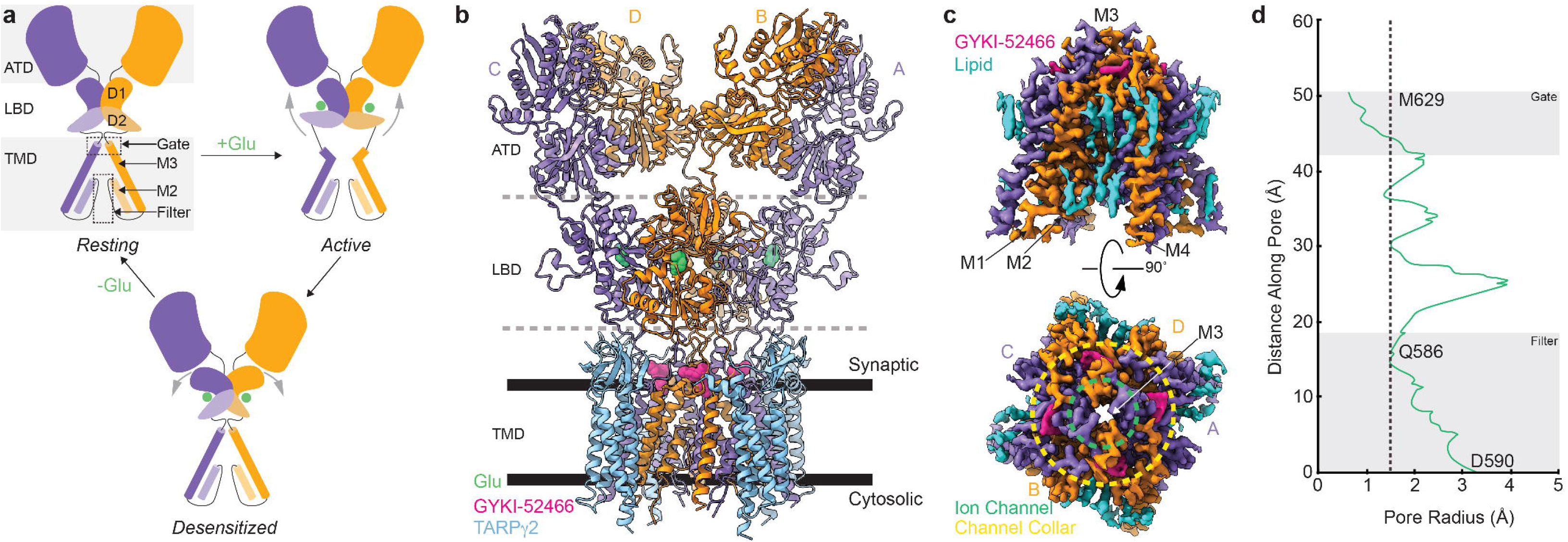
Structure of the AMPAR inhibited state. a) Schematic representation of the AMPAR gating cycle. b) Ribbon illustration detailing the structure of the AMPAR inhibited state, IS-1. GluA2 subunits are purple (A/C) or orange (B/D) depending on their positions. GYKI- 52466 (pink) is bound at all four TMD collar regions and each LBD clamshell is closed around glutamate (green). TARPγ2 subunits (light blue) occupy all four auxiliary sites around the receptor. c) High resolution details of the focused GluA2 TMD from cryo-EM reconstruction. Side view (top) of the GluA2 TMD from showing the M3 bundle crossing in a closed conformation and top view (bottom) showing the bundle crossing constricting access to the ion channel (green, dashed) and the relative location of the channel collar (yellow, dashed) with GYKI-52466 bound to all four GluA2 subunits. Lipids (blue) adorn the AMPAR TMD. d) Plot of the ion channel radius along the pore axis showing a constriction at the M3 bundle crossing gate. Dashed line represents the radius of a water molecule.

Allosteric modulators bind to AMPARs at sites distinct from the Glu binding site in the LBD and bias AMPAR function. Positive allosteric modulators such as cyclothiazide (CTZ) and its derivatives bind between the D1’s of local LBD dimers and enhance D1-D1 contact during activation, thus favoring activation and preventing AMPAR desensitization^19–22^. How negative allosteric modulators such as PPLMs prevent AMPAR activation is more enigmatic. The current paradigm in the field is that PPLMs bind to the region of the TMD that is extracellular facing and prevent AMPARs from transitioning to the active state^4,9^. How this occurs is unknown because AMPARs have not been studied structurally in the presence of both Glu and PPLMs^9,23^. Thus, while these studies describe binding sites, the mechanism of inhibition is unclear because the PPLMs are bound to resting state receptors.

We hypothesized that PPLMs allosterically inhibit AMPARs by pushing the receptors into a desensitized-like state that decouples Glu binding from the ion channel. Furthermore, we expected there to be allosteric competition between positive modulators such as CTZ and negative modulators such as PPLMs to control AMPAR function because their binding sites in AMPARs are at distinct sites. How this would occur is unknown and not yet observed in AMPARs nor any family of ligand gated ion channels.

To test these ideas, we activated AMPARs in the presence of both the negative allosteric modulator GYKI-52466 and positive allosteric modulator CTZ. Through a combination of cryo-electron microscopy (cryo-EM), single-molecule fluorescence resonance energy transfer (smFRET), electrophysiology, and molecular dynamics simulations, we demonstrate that GYKI- 52466 binding in the TMD decouples Glu binding from the ion channel by allosterically rearranging the AMPAR LBD. In addition, GYKI-52466 binding in the TMD prevents positive allosteric modulation by CTZ in the LBD. Our findings shed new insights into how allosteric modulation is coordinated across AMPARs, demonstrate allosteric competition between modulators, and will invigorate structure-based drug design against AMPARs and provide a framework for studying inhibition across iGluRs and other ligand-gated ion channels.

## Results

### Structures of inhibited AMPAR complexes

To identify how PPLMs inhibit AMPARs via negative allosteric modulation and test our idea of allosteric competition, we purified homotetrameric AMPARs composed of the GluA2 subunit in complex with the auxiliary subunit transmembrane AMPAR regulator protein (TARP)γ2^15,18,24^, which enhances AMPAR activation (**Extended Data Fig. 1a,b**), and preincubated the receptors with cyclothiazide (CTZ), a positive allosteric modulator that inhibits AMPAR desensitization and allows the activated state of the receptor to be captured with cryo-EM^15,25^. We activated these AMPAR complexes in the presence of GYKI-52466 to capture inhibited states (**Extended Data Fig. 1c**). We captured inhibited states through two different schemes. In the first scheme (inhibited state 1, IS-1), we mixed the CTZ-bound receptors with Glu and GYKI-52466 immediately prior to freezing. In the second scheme (IS-2), the receptors were pre-incubated with GYKI-52466 in addition to CTZ, and Glu was added immediately before freezing. Each approach resulted in similar inhibited states (**Extended Data Fig. 2a**).

We focus our analysis on IS-1 owing to higher data quality (**Extended Data Figs. 3,4**). The overall structure of the AMPAR complexes reveal key details of an inhibited AMPAR (**Fig. 1b**). There is an overall “Y” arrangement of the receptor, with the two-layered ECD comprised of the ATD and LBD. All four GluA2 LBDs are Glu-bound. Immediately below is the GluA2 TMD, which is fully occupied with four TARPγ2 auxiliary subunits. Four GYKI-52466 molecules are bound to the TMD along its extracellular-facing surface.

Cryo-EM reconstruction of the AMPAR TMD to 2.6 Å enables elucidation of key features of the AMPAR TMD during inhibition. The four GYKI-52466 molecules are wedged between helices at the top of the TMD (**Fig. 1c**). Importantly, the GYKI-52466 binding sites are adjacent to the ion channel in the channel collar region. The collar channel forms a ring of solvent accessible pockets for PPLMs that surrounds the M3 gate. Lipids adorn the AMPAR TMD on both the extracellular and cytosolic facing (**Fig. 1c**) and are critical to plug cavities within the bilayer that would otherwise perturb the solvent accessibility of the ion channel (**Extended Data Fig. 5**). Next, we measured the ion channel radius, which indicates a closed channel; the upper channel gate, defined by M629 at the M3 helix crossing, completely restricts channel access (< 1.0 Å radius) to both water molecules and sodium ions (**Fig. 1d**).

While both activator (Glu) and negative allosteric modulator (GYKI-52466) are bound to the AMPAR, the positive allosteric modulator CTZ, which was included in the preparation of both IS- 1 and IS-2, is absent from both cryo-EM reconstructions. This indicates that the states we captured are markedly different from previously captured states of AMPARs, as CTZ binds to both the resting and activated states of the receptor^25–27^. Thus, GYKI-52466, at a binding site completely distinct from that of CTZ in the AMPAR LBD allosterically outcompetes CTZ.

### The GYKI-52466 binding site

Reconstruction of the AMPAR TMD enabled precise building of the AMPAR TMD (**Extended Data Fig. 6a**). However, to delineate the precise details of GYKI-52466 binding in the AMPAR channel collar we used symmetry expansion to reconstruct the binding site to 2.2 Å resolution (**Extended Data Fig. 6b-e**). This enabled us to characterize the complete binding pocket (**Fig. 2a,b**). GYKI-52466 is partially stabilized in the collar through a π-bond stack where GYKI-52466 is sandwiched between F623 at the top of the M3 helix and the P520 on the pre-M1 helix (**Fig. 2a**). Van der Waals forces from five nearby residues, S516, N619, S615, Y616, and N791 also contribute to the binding site (**Fig. 2b**). The N3 of GYKI-52466 is sandwiched between both Y616 on M3 and S615 on the M3 of an adjacent subunit. Therefore, GYKI-54266 is wedged between two AMPAR subunits in the TMD (**Fig. 2b**). Unique to GYKI-52466 pocket from other PPLM-bound structures^9,23^ is the likely contribution of water molecules to the binding pocket (**Extended Data Fig. 6e**) and coordination by N791 on M3 (**Fig. 2a**).

**Figure 2.**
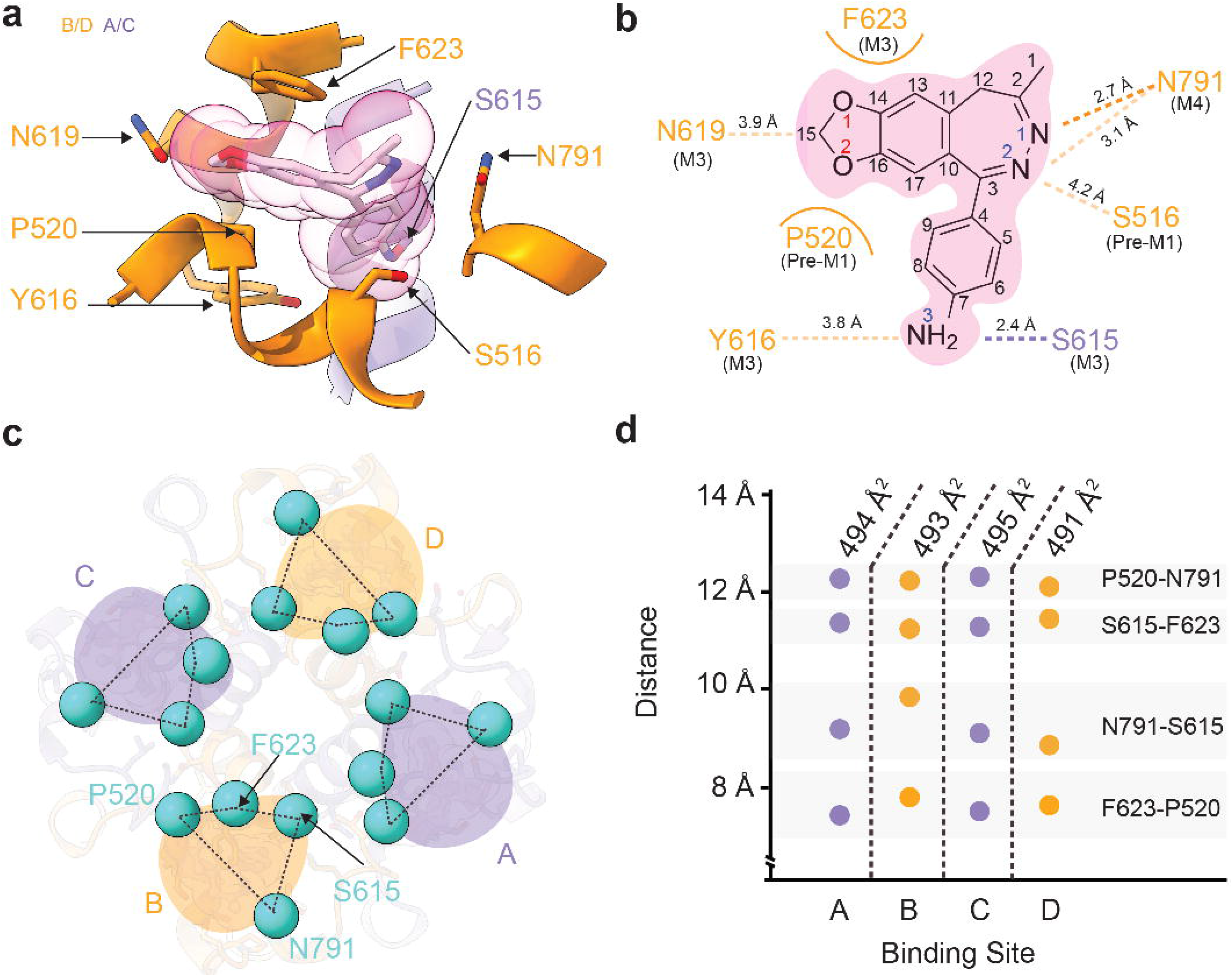
High-resolution details of GYKI-52466 binding. a) GYKI-52466 (gray with pink spheres) makes extensive contact with residues within the TMD collar region (orange ribbons), including interacting with S615 on the neighboring M4 helix counterclockwise from the ‘bound’ subunit (purple). b) Schematic representation of GYKI-52466 interactions with TMD collar residues. Pi bonds represented by curved lines; Van der Waals interactions represented by dashes. c) Top-down view of the inhibited TMD, showing landmark residues around the GYKI- 52466 binding pocket. d) Plot detailing the inter-residue distances between landmark GYKI- 52466 binding pocket residues and solvent accessible surface surrounding GYKI-52466.

During AMPAR activation the AMPAR subunits in the B and D positions undergo the most dramatic conformational changes in the TMD to drive opening of the ion channel.^15,16,28^ Because of this, we hypothesized that the binding pocket around GYKI-52466 may be more compact in the B/D positions during inhibition because GYKI-52466 may directly block the conformational changes in the B/D positions that are associated with activation.

To assess the overall shape and size of the binding pockets in each subunit, we measured the distances between P520, N791, S615 and F623 (**Fig. 2c**). To our surprise, the binding pockets were remarkably similar as indicated by inter-residue distances (**Fig. 2d**). On average, there is a ∼12 Å distance between pairs P520-N791 as well as S615-F625, ∼9Å distance between N791- S615, and ∼8 Å distance between F623-P520. Thus, the shape around the GYKI-52466 binding site is roughly the same in each binding pocket. The solvent accessible surface around GKYI- 52466 is also similar in each AMPAR subunit position, with an average solvent accessible surface of ∼493 Å^2^ around GYKI-52466. Thus, despite activation being accompanied by dramatic rearrangements in the B and D subunits to accommodate ion channel opening, there are no discernible differences between subunit positions in the inhibited state. Thus, we expected that the mechanism of allosteric inhibition by GYKI-52466 is driven through an alternative mechanism, despite the GYKI-52466 binding site being within the TMD.

### GYKI-52466 Decouples Ligand Binding from Ion Channel Opening

To elucidate the inhibition mechanism, we compared our inhibited state structure to an activated state AMPAR structure (**Fig. 3a**). The majority of the TMD is largely similar between the two states, except at the channel gate, which is formed by the top of the M3 helices (**Fig. 3a**). During activation, the M3 helices kink outward from the pore axis to open the channel. This key movement is blocked by the presence of GYKI-52466 in the B/D AMPAR subunit positions due to the presence of GYKI-52466 in the channel collar (**Fig. 3a**, inset i). However, there are no key differences between the GYKI-52466 B/D and A/C positions of the channel collar in the inhibited state (**Fig. 2d**). In addition, each individual LBD in the inhibited state is Glu-bound, with a similar overall conformation to individual LBDs in the activated state (**Fig. 3b**). We hypothesized that the conformational changes that dictate AMPAR inhibition occur in the LBD layer because the ATD does not play a significant role in gating^29–31^.

**Figure 3.**
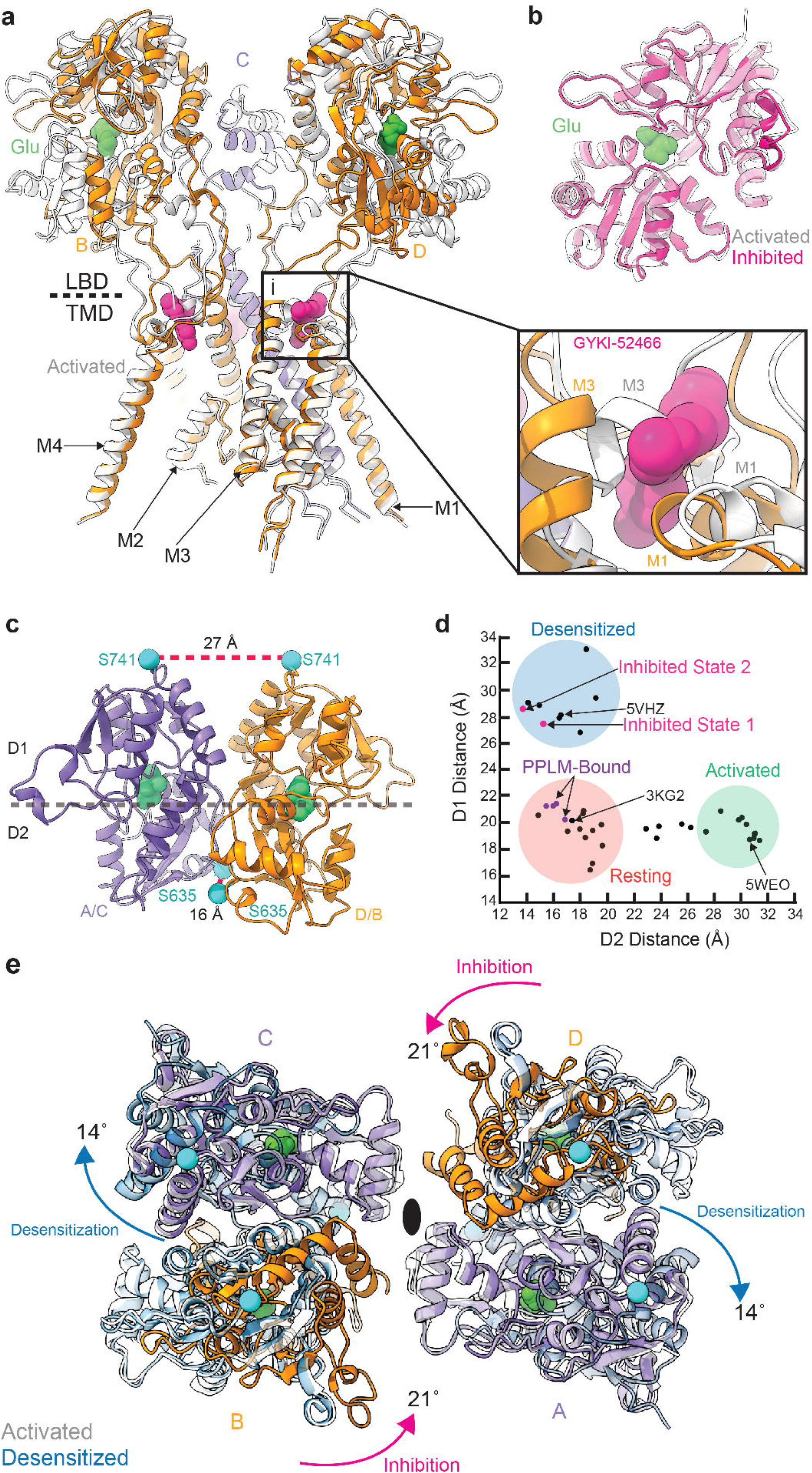
Mechanism of negative allosteric modulation. a) Overlay of the inhibited state (orange) and the activated state (pdb 5WEO, white). *Inset:* Close-up view of the GYKI-52466 binding site revealing a steric clash with the kinked M3 helix found in the open state. b) Overlay of isolated LBD clamshells from the inhibited state (pink) and the open state (white). c) Local clamshell dimers within the LBD layer viewed from behind, showing the relative distances of the D1 and D2 lobes of the LBD dimer as illustrated by the landmark residues S741 (D1) and S635 (D2). d) Plot of the D1 distance (Cα of S741) versus D2 distance (Cα of S635) measured for representative AMPAR structures captured in either the resting, activated, or desensitized state. Based on these measurements, the inhibited states (pink) cluster most closely with the desensitized state structures. e) Overlay of the overall LBD layer of the inhibited state (orange and purple), activated state (white) and desensitized state (blue) viewed from the top or extracellular side. Desensitization causes a 14° clockwise rotation of the A/C subunits within the LBD layer relative to the activated state. In contrast, inhibition drives a counterclockwise rotation of the B/D subunits within the LBD layer relative to the active state.

The inhibited AMPAR LBD layer is markedly different than the activated state (**Fig. 3a**). While the individual LBDs in each protomer share the same Glu-bound conformation (**Fig. 3b**), the LBD dimers undergo a significant conformational change to accommodate AMPAR inhibition. To assess the changes, we measured the distances between the D1-D1 and D2-D2 in LBD local dimers, which are major indicators of the functional state of the AMPAR^29^. For example, during activation, the distances between D1’s in LBD local dimers are decreased as the D2’s separate to pull open the ion channel (**Fig. 1a**). During desensitization, the opposite occurs, where the D1’s separate, and the D2 interface is minimized, which decouples Glu binding from the channel, which allows it to close (**Fig. 1a**). The D1 and D2 interfaces in the resting state of the receptor represent an intermediate between the extremes of activation and desensitization.

In IS-1, we measured the distances between the Cα atoms of S741 (D1 separation) and between the Cα atoms of S635 (D2 separation) (**Fig. 3c**). The D1 interface is markedly separated (27 Å) compared to the D2 interface (16 Å). To assess how these separations fit with the conformational landscape of existing AMPAR structures in the protein data bank (pdb), we measured D1 and D2 separation in AMPAR structures deposited in the pdb (**Fig. 3d**). Generally, structures with a ∼26 Å or greater distance between S741 in D1 residues represent a desensitized state, while structures with a ∼27 Å or greater distance between S635 in D2 represent an active state, with resting state structures representing a medium between the two separations. The activated state of AMPAR is exemplified by pdb 5WEO, resting state pdb 3KG2, and desensitized state pdb 5VHZ (all pdbs are mapped in **Extended Data Fig. 7**). The significant rupturing of the D1 interfaces in both IS-1 and IS-2 places these LBD dimers squarely into the desensitized classification of LBD dimers. Critically, existing PPLM-bound structures in the pdb represent the resting state of the receptor because they are not Glu-bound (**Fig. 3d**). This is marked by significant differences across the receptors between the PPLM-bound apo states and the inhibited states from this study (**Extended Data Fig. 2b**).

While the LBDs in local dimers are in a desensitized-like state, the total motion of the LBD layer reveals that inhibition is unique from desensitization. During desensitization, the A/C subunits roll away from their B/D partners to facilitate separation within the local dimers and decouple Glu binding from the ion channel^18,32^ (**Fig. 3e**). In inhibition, we observe the opposite (**Fig. 3e**). During inhibition, the B/D LBDs rotate 21˚ counterclockwise away from their A/C counterparts, which appear to maintain the position that they assume in the active state (**Fig. 3e**). Therefore, like their role in activation, the B/D subunits drive inhibition. We expect that because the M3 helix kink is prevented by GYKI-52466 in the B/D subunits (**Fig. 3a**), this drives rearrangement in the LBD by the same subunits to accommodate inhibition.

### Allosteric Competition to Control the AMPAR LBD

The positive allosteric modulator CTZ is absent in both our inhibited-state cryo-EM reconstructions despite being included during preparation. This suggests that GYKI-52466, a negative allosteric modulator that binds in the AMPAR TMD, outcompetes CTZ, which binds between AMPAR LBDs, to allosterically control AMPAR function. The absence of CTZ in our cryo-EM reconstructions may be explained by the separation of the D1-D1 interface that we observe in the inhibited state (**Fig. 4a**), which effectively ruptures the CTZ binding pocket (**Fig. 4b**). CTZ binding between AMPAR LBDs at the D1-D1 interface stabilizes D2-D2 separation and prevents desensitization. However, inhibition by GYKI-52466 prevents CTZ from stabilizing the D1-D1 interface as the CTZ binding site is completely ruptured in inhibition.

**Figure 4.**
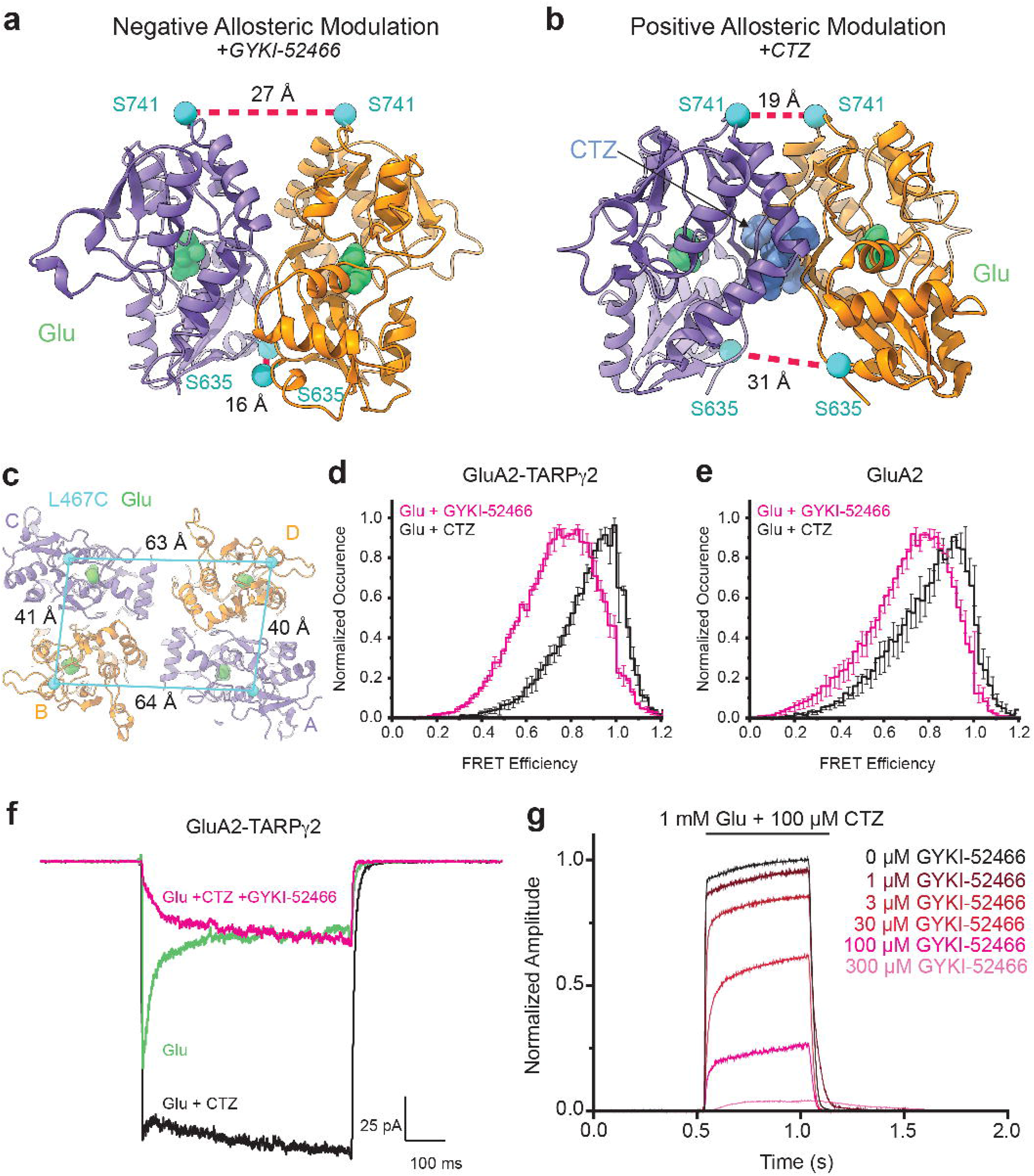
Allosteric Competition in AMPARs. a) Local LBD clamshell from the inhibited state (IS-1) with landmark residues to show D1 and D2 separation within the dimer. This represents the dimer during negative allosteric modulation. b) Local LBD clamshell dimer activated in the presence of CTZ, showing decreased D1 separation and increased D2 separation. This is indicative of positive allosteric modulation. CTZ shown in blue. Distances measured as in Figure 3. c) Top-down view of the LBD layer with L467, where the L467C mutation is used for maleimide dye labeling, marked with a blue sphere and inter-subunit L467-L467 Cα distances labeled. d) Plot of FRET efficiency between LBD clamshells within a local dimer when GluA2-TARPγ2 protein was treated with Glu + CTZ alone vs Glu + CTZ + GYKI-52466. e) Same as in *d*, but with GluA2 protein in the absence of TARPγ2. f) Whole-cell patch clamp recordings from HEK293 cells expressing GluA2-TARPγ2 protein in the presence of either Glu, Glu + CTZ, or Glu + CTZ +G YKI-52466. g) Whole-cell patch clamp traces of HEK293 cells expressing GluA2-TARPγ2 treated with 1mM Glu + 100 µM CTZ either alone or with increasing concentrations of GYKI-52466.

We therefore postulated that the negative allosteric modulator GYKI-52466 and the positive allosteric modulator CTZ might compete for influence over AMPAR gating, despite binding at disparate sites, with GYKI-52466 exerting greater influence than CTZ when both are at saturating concentration in the preparation. We refer to such a paradigm as allosteric competition.

To directly test the effects of negative and positive allosteric modulation on the D1-D1 interface, we directly assayed the separation of the D1-D1 interface with smFRET. We achieved this with the mutation L467C at the top of D1 to enable attachment of a dye by maleamide chemistry and establish FRET pairs at the top of the GluA2 LBD^33^ (**Fig. 4c**). Labeling GluA2 homotetrameric AMPARs at L467C is ideal for measuring distances within LBD dimers; the FRET efficiency when GluA2 is in the activated state (Glu + CTZ) is expected to be ∼92% within an LBD dimer and ∼19% across dimer pairs when Alexa-555 and Alexa-647 are used as the donor-acceptor pair^34^.

We tested coupling of the D1 interface in our GluA2-TARPγ2 construct during positive allosteric modulation in the presence of both 1 mM Glu and 100 µM CTZ (**Fig. 4d**), where the D1s between LBD dimer pairs are at their closest^34–36^ (**Fig. 4b**). The Glu and CTZ smFRET efficiency histogram shows higher efficiency than receptors in negative allosteric modulation (1 mM Glu and 100 µM GYKI-52466; **Fig. 4d**). This indicates that the distance across the D1 interface is shorter in the presence of the positive modulator CTZ than in the presence of the negative modulator GYKI- 52466. To test that the decrease in smFRET efficiency in inhibitory conditions is AMPAR-dependent and not TARP-dependent, we also tested smFRET efficiency in GluA2 homotetramers in the absence of TARPγ2 (**Fig. 4e**). A similar effect is seen between positive allosteric (Glu + CTZ) and negative allosteric (Glu + GYKI-52466) conditions with GluA2 alone, which points to the decrease in smFRET efficiency not being TARP-dependent but GYKI-52466 or CTZ dependent.

The individual smFRET traces show that the protein occupies 2-3 FRET efficiency states (**Extended Data Fig. 8**) with the most probable state having a FRET efficiency of 0.93 in the presence of cyclothiazide and 0.82 in the presence of GYKI (**Extended Data Fig. 8**). These FRET efficiencies correspond to distances of 33 Å and 39 Å, respectively. The distance change of 6 Å agrees with our IS-1 and IS-2 cryo-EM structures which show a D1-D1 (L467) distance change of 6 Å when compared to the CTZ-bound, activate state AMPAR structure^15^. Thus, separation of the D1s in AMPAR LBD dimers appears to be due to negative allosteric modulation by GYKI- 52466.

Collectively, our observations from smFRET and cryo-EM suggest that negative and positive allosteric modulation occupy dramatically different conformational states. The differences between the conformational spaces in each state are a potential mechanism for allosteric competition between the two modulators (**Fig. 4a,b**). Thus, we hypothesized that the negative allosteric effects of GYKI-52466 would outcompete CTZ’s positive allosteric modulation on AMPARs. This would be reflected in GYKI-52466 negating CTZ’s non-desensitizing effects that CTZ has on AMPAR currents.

We tested whether GYKI-52466 could overcome the positive allosteric effect CTZ has on AMPARs by patch clamp electrophysiology of HEK293 cells expressing GluA2-TARPγ2 (**Fig. 4f**). In the absence of CTZ, the currents rapidly desensitize when treated with 1 mM Glu, and desensitization is completely ablated with 100 µM CTZ. However, the effect of CTZ on desensitization is negated by 100 µM GYKI-52466. Thus, the inhibitory effect of GYKI-52466 on AMPARs outcompetes the positive effect that CTZ has on AMPAR currents. We complemented the observation that GYKI-52466 has greater control on AMPAR allostery than CTZ with dose-response curves (**Fig. 4g**). This data shows that GYKI inhibits GluA2-mediated currents even in the presence of excess CTZ; at ∼3x CTZ concentration, GYKI-52466 inhibits 50% of AMPAR currents (**Fig. 4g**). The IC_50_ of GYKI-52466 in the presence of CTZ was determined to be 39.87 ± 6.75 μM (p = .00022). This is a ∼10x reduction of the IC_50_ compared to GYKI-52466 alone on AMPAR-TARP complexes^37^, which aligns well with the observed 10x reduction in GYKI-52466 IC_50_ on AMPARs in the presence of CTZ^26,38,39^.

While our electrophysiology data agrees with previously reported observations, our data in the context of our observations from cryo-EM and smFRET demonstrates that GYKI-52466 binding in the TMD outcompetes CTZ binding in the LBD for allosteric control of AMPARs.

### Free Energy Landscape Governing Rupture of the LBD Dimer Interface

Comparison of singular inter-residue distances may not account for how inhibition and desensitization occupy similar conformations within LBD dimers but an overall distinct conformation in the overall LBD tetramer^27,40–43^ (**Fig. 3e**). We hypothesized that this may be due to distinct free energy minima accompanying inhibition and desensitization. To test this, we computed a two-dimensional free energy landscape, or potential of mean force (PMF), governing rupture of a Glu-bound GluA2 LBD dimer interface using umbrella sampling free energy molecular dynamics (MD) simulations.

Our PMF is a function of a two-dimensional order parameter (χ_1_ and χ_2_) that reports global changes within an LBD dimer. χ_1_ and χ_2_ describe the distances between the center of mass of helix J in D1 and the center of mass of helix D in D1 on a partner LBD in the dimer (**Fig. 5a**). (χ_1_, χ_2_) differs from the one-dimensional collective variable previously used by to examine LBD dimer stabilities in AMPARs and kainate receptors via steered MD simulations^44^. While the LBDs are generally symmetric, the order parameter is not (**Fig. 5a**); χ_1_ describes the helix J and D distance that is exterior facing, while in the context of a tetramer, χ_2_ describes the helix pair that faces the interior of the AMPAR. Thus, this enables a two-dimensional approach to characterizing global changes in the LBD dimers.

**Figure 5.**
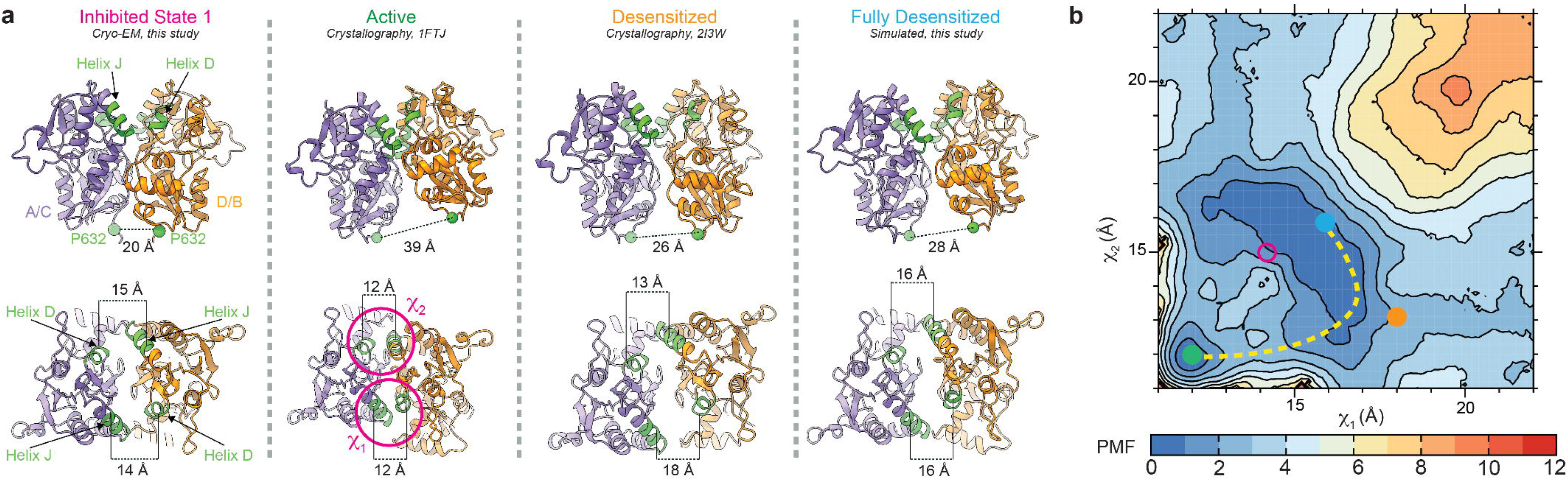
Free energy landscapes governing desensitization and inhibition. a) (top) Rear views of local LBD clamshell dimers from the inhibited state, activated state, desensitized state, or a projected maximally desensitized state. (bottom) Top-down views of the inhibited, activated, desensitized, or maximally desensitized states with Helices D and J, colored green, labeled in accordance with their contribution to the 2D order parameter. b) Free energy landscape governing LBD dimer conformations, i.e., the separation of helices D and J at the dimer interface, with the measured distances of the activated (green), desensitized (orange), and maximally desensitized distances (blue) plotted on the diagram. Corresponding measurements from the inhibited state are shown in pink, which corresponds to where we would expect the inhibited state to sit within the free energy landscape. The dashed line suggests the most probable transition pathway between the active and desensitized conformations. The free energy landscape is contoured in 1 kcal/mol increments.

Through sampling χ_1_ and χ_2_ in the context of Glu-bound LBDs, we can understand the energetics associated with rupturing the D1-D1 interface. Conformers for the umbrella sampling windows were generated using targeted MD simulations initiated from the crystal structure of an activated GluA2 LBD and using the crystal structure of a desensitized GluA2 LBD as a guide (**Fig. 5a**; see Methods)^45,46^. Sampling windows were established in 1 Å increments along χ_1_ and χ_2_. The activated state LBD dimer occupies a small free energy basin within the PMF, whereas the fully desensitized LBD occupies a significantly larger basin (**Fig. 5b**). The crystallized desensitized LBD, stabilized by a disulfide bond, lies near the most probable transition pathway between the active and desensitized conformations. This pathway suggests that during rupture of the dimer interface, one J-D helix pair breaks before the other rather than both pairs breaking simultaneously, thereby circumventing a free energy barrier separating the two basins. The broader free energy basin associated with desensitization compared with activation may account for how short-lived the active state is compared to the longer-lived desensitized state.

A point mutation, L483Y in helix D, had been identified to strongly stabilize the non-desensitized (active) state^47^. To test whether our umbrella sampling strategy could recapitulate the effect of this mutation, we performed an analogous free energy calculation using the GluA2-L483Y LBD dimer. Umbrella sampling window conformers were generated from the crystal structure of the L483Y LBD dimer^22^. The PMF of this non-desensitizing mutant reveals a substantially reduced free energy basin for the desensitized state, transforming the active state basin into the global free energy minimum (**Extended Data Fig. 9**).

In inhibition, we observe separation of χ_1_ and χ_2_ compared to the activated LBD dimer (**Fig. 5a**). Interestingly, this likely occupies a PMF basin that is distinct from the pathway of desensitization (**Fig. 5b**). This supports the observation that inhibition is similar, but distinct, from desensitization. The two-dimensional order parameter that we sample in this experiment accounts for how the LBDs within a dimer pivot away from each other to accommodate D1 separation. We hypothesize that the distinct free energy basins of inhibited and desensitized LBDs account for the differences we observe in the overall motion in the LBD tetramer during inhibition and desensitization (**Fig. 3e**).

## Discussion

Allosteric modulation of AMPARs is a critical avenue for small molecule therapeutics to regulate AMPAR function. While mechanisms for positive allosteric modulation have been determined prior to this work, the mechanisms of allosteric inhibition by negative allosteric modulators have remained unknown. In addition, we outline how negative and positive modulators can compete in what we term allosteric competition. To our knowledge, this is the first description of allosteric competition in AMPARs, iGluRs, and ligand gated ion channels. We delineate the principles of allosteric competition and negative allosteric modulation with cryo-EM, smFRET, electrophysiology, and molecular dynamics. Elucidation of these details complete our knowledge of the allosteric landscape and provide foundations for structure-based drug design.

Indeed, despite the central role of AMPARs in neuronal signaling and their role in disease, therapeutic development against AMPARs has remained challenging. This is reflected by only a single AMPAR-targeting drug, perampanel, being approved in the United States and Europe. Over 400,000 people have been prescribed this drug across all indications^48^. Despite this, the perampanel mechanism of action has remained unknown until this study. Perampanel, like GYKI- 52466, and other PPLMs, bind to AMPARs in the ion channel collar region and inhibit AMPARs. Our data show that noncompetitive inhibitors do not simply inhibit the activation of AMPARs by keeping the receptor in the resting state^9,26^ but instead inhibit the receptor by decoupling Glu binding from channel opening by pushing the receptor into a desensitized-like state (**Fig. 6a**). The inhibited states (IS-1 and IS-2, this study) show marked differences compared to AMPAR structures bound to PPLMs in the resting state (**Extended Data Fig. 2b**). There are major differences in the separation of the D1-D1 interface (**Fig. 3d**) and overall LBD layer, though the TMD and ATD are similar (**Extended Data Fig. 2b**).

**Figure 6.**
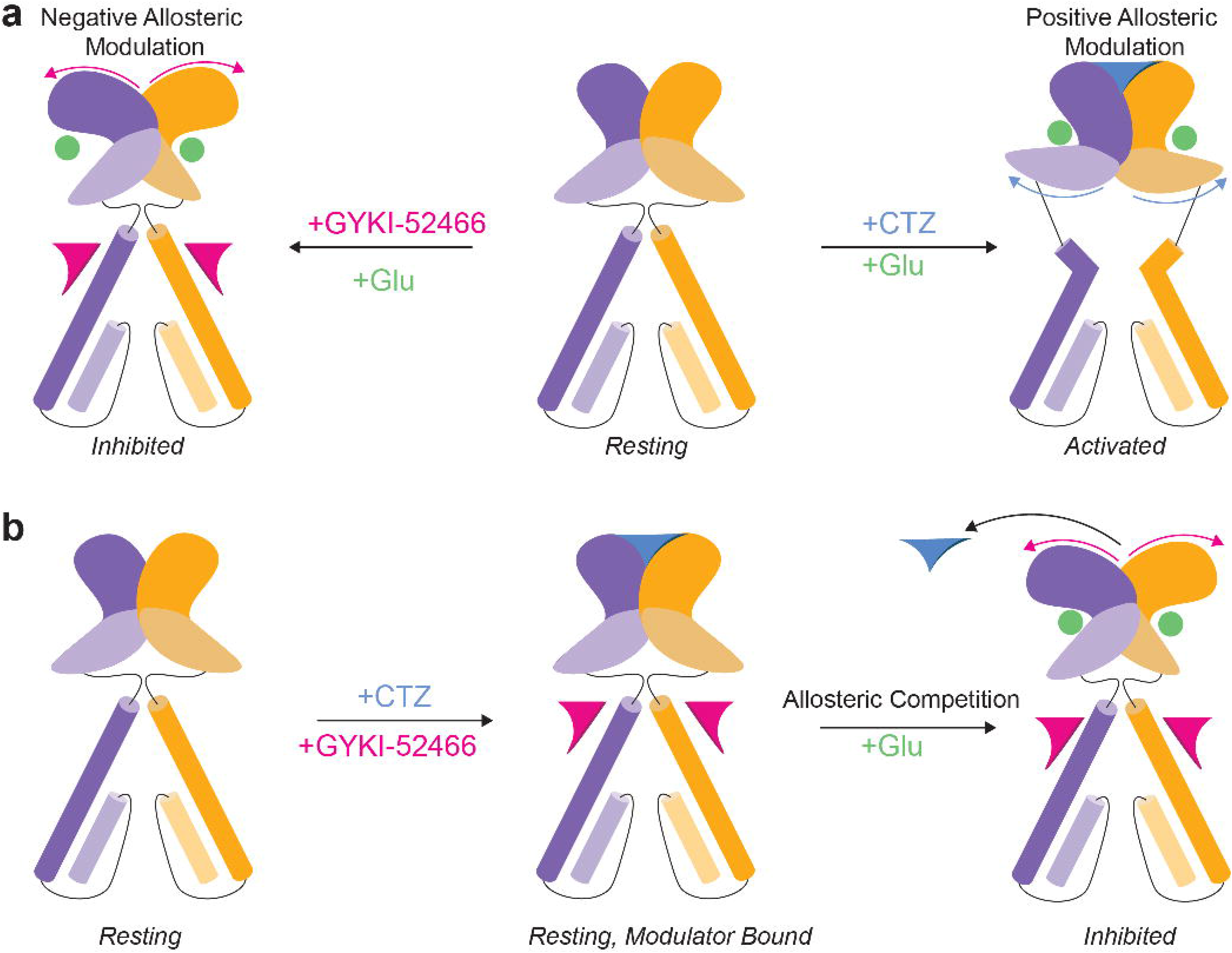
Allosteric landscape of AMPARs. a) Activation of AMPARS in the presence of the negative allosteric modulator GYKI-52466 produces the inhibited state, which we show in this study. In contrast, activation of AMPARs in the presence of the positive allosteric modulator CTZ produces the activated state. b) When both CTZ and GYKI-52466 are present, both can bind to the resting state of the receptor, but after Glu binding, GYKI-52466 outcompetes CTZ to control the AMPAR LBD, resulting in CTZ being displaced, and inhibition of the receptor.

Supporting our inhibition mechanism are previous studies that suggested a two-step mechanism of inhibition, where an initial binding event by PPLMs is insufficient to produce complete inhibition^49,50^. These findings support our results, where a two-step mechanism would constitute GYKI-52466 binding followed by Glu binding in the LBDs and rupturing of the D1 interface within LBD dimers. This demonstrates how binding of PPLMs in the ion channel collar allosterically controls the AMPAR ECD (**Fig. 6a**). The evidence for inhibition occupying a similar but distinct mechanism to desensitization also provides new data to conceptualize therapeutically targeting AMPARs. For example, the B/D subunit position domain motion that accompanies inhibition (**Fig. 3e**) may provide a route for specificity in small molecule targeting considering that these positions are enriched for specific GluA subunits in native AMPARs^51–53^.

Our structural data predict that the mechanism of inhibition by PPLMs is independent of auxiliary subunit occupancy of AMPARs. This prediction agrees with our smFRET data showing similar shifts in FRET efficiency in the presence or absence of TARPγ2 in the AMPAR (**Fig. 4b**). However, noncompetitive inhibition of AMPARs may function similarly across different drug types. AMPARs are tightly regulated by TARPs and other auxiliary subunits^23,24,28,32,54–56^ and recently identified compounds demonstrate selectivity for particular AMPAR-TARP complexes^57–62^. Structural investigations of the binding sites reveal that drug binding of TARP-dependent inhibitors is at a site distinct from that of PPLMs; the site is within the interface between TARPs and AMPAR transmembrane helices. It is possible that these TARP-dependent noncompetitive inhibitors also act as negative allosteric modulators by forcing receptors into a desensitized-like state to achieve inhibition, despite having binding sites distinct from PPLMs. However, resolving this question will require additional studies with AMPARs activated in the presence of the TARP-dependent noncompetitive inhibitors.

A key observation from our data is the competing mechanisms of positive allosteric modulators (e.g., CTZ) and negative allosteric modulators (e.g., PPLMs). CTZ positively modulates AMPAR function by bringing together the D1-D1 interface in LBD dimers (**Fig. 6a**). Thus, GYKI-52466 and CTZ have been consistently reported to produce opposing effects on channel conductance^9–11,63^. Early studies postulated that GYKI-52466 and CTZ bind to the same site on AMPARs, owing to their countervailing effects on AMPAR channel conductance^63,64^. However, subsequent structural and physiological data revealed that CTZ and PPLMs act at distinct sites^9,26,65^, thereby rendering their mechanistic competition unclear.

Our data indicate that negative allosteric modulators such as GYKI-52466 can outcompete positive allosteric modulators that bind to a completely distinct site such as CTZ. Indeed, both PPLMs and CTZ can bind to resting state AMPARs^9,15,23,25,49,66,67^. This allosteric competition (**Fig. 6b**) to control AMPAR function is a new way to understanding allosteric modulation and allosteric competition in ligand-gated ion channels. This idea was alluded to in early studies on the interplay between GYKI-52466 and CTZ^26,39^. It was suggested that there is a negative allosteric effect from GYKI-52466 that does not prevent CTZ binding, but instead alters the effect of CTZ on AMPAR desensitization. Our work shows that negative allosteric modulation by GYKI-52466 prevents CTZ from positively modulating AMPARs through rupturing the CTZ binding site. The competing mechanisms of PPLMs such as GYKI-52466 and CTZ may provide a route to fine tune the behavior of AMPARs^11^.

Our data reveal that inhibition is a distinct state from the resting state (**Fig. 3e**), but because individual inhibited channels bypass the open state entirely to arrive at the inhibited state, no current is passed and therefore this state is not detectable with electrophysiological analysis other than a reduction in peak current amplitude in whole cell recordings. Thus, our proposed mechanism bridges electrophysiological studies of GYKI-52466 and PPLM competition with CTZ with the structural identification of PPLM binding sites. Furthermore, the revealed details of the GYKI-52466 binding site allow us to disambiguate the arrangement and contribution of sidechains in the PPLM binding pocket that were previously described^9^. Consistent with mutagenesis studies conducted in the PPLM binding pocket^9^, we can confirm the direct interaction of key sidechains that reduced inhibition efficiency following mutation, including S516, P520, S615, F623 and N791 as well the likely involvement of N619 in stabilizing GYKI-52466 specifically. Importantly, water molecules may play an important role in the coordination of small molecules in this binding pocket (**Extended Data Fig. 6e**). While the residues that coordinate GYKI-52466 are largely conserved across AMPAR subunits (**Extended Data Fig. 10**), the high-resolution details outlined here, and identification of the negative allosteric modulation mechanism, will improve small molecule design in future studies.

In sum, we reveal the mechanism of action of how AMPARs are allosterically inhibited and how a new phenomenon of allosteric competition can occur within AMPARs. Our data provide foundations for structure-based drug design against AMPARs as well as a framework to study the mechanism of action of noncompetitive inhibition in other ligand-gated ion channels.

---

**Extended Data 1.**
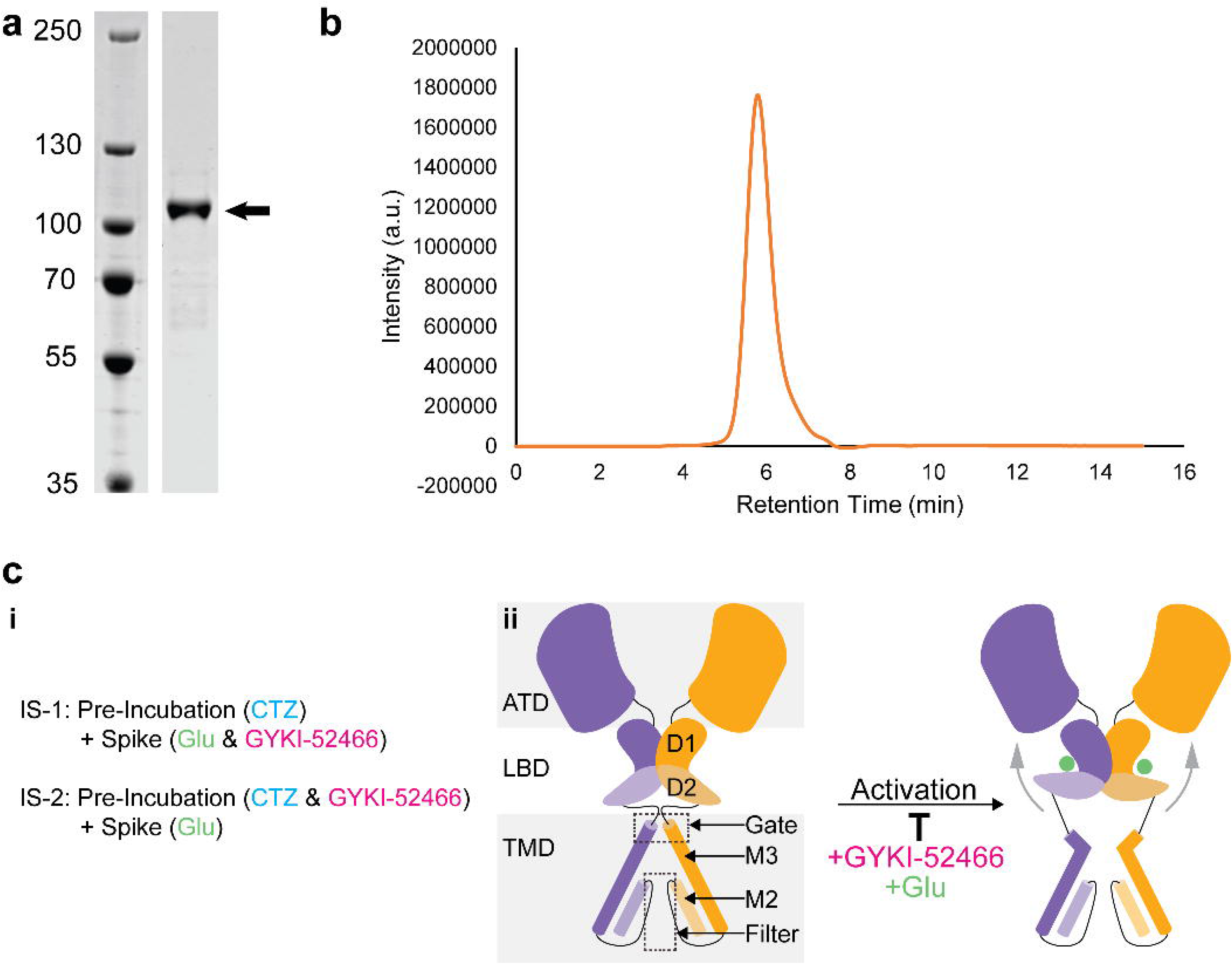
Preparation of GluA2-TARPγ2. a) Coomassie-stained SDS PAGE gel of purified GluA2-TARPγ2 sample showing a single band at the predicted molecular weight (arrow). b) Size exclusion chromatogram of purified GluA2-TARPγ2 sample showing a single monodispersed peak at the predicted retention time for a GluA2-TARPγ2. c) (i) Treatment regimens for producing the different inhibited states IS-1 and IS-2, (ii) cartoon demonstrating the targeted outcome of activating the GluA2-TARPγ2 assembly in the presence of the inhibitor GYKI-52466.

**Extended Data 2.**
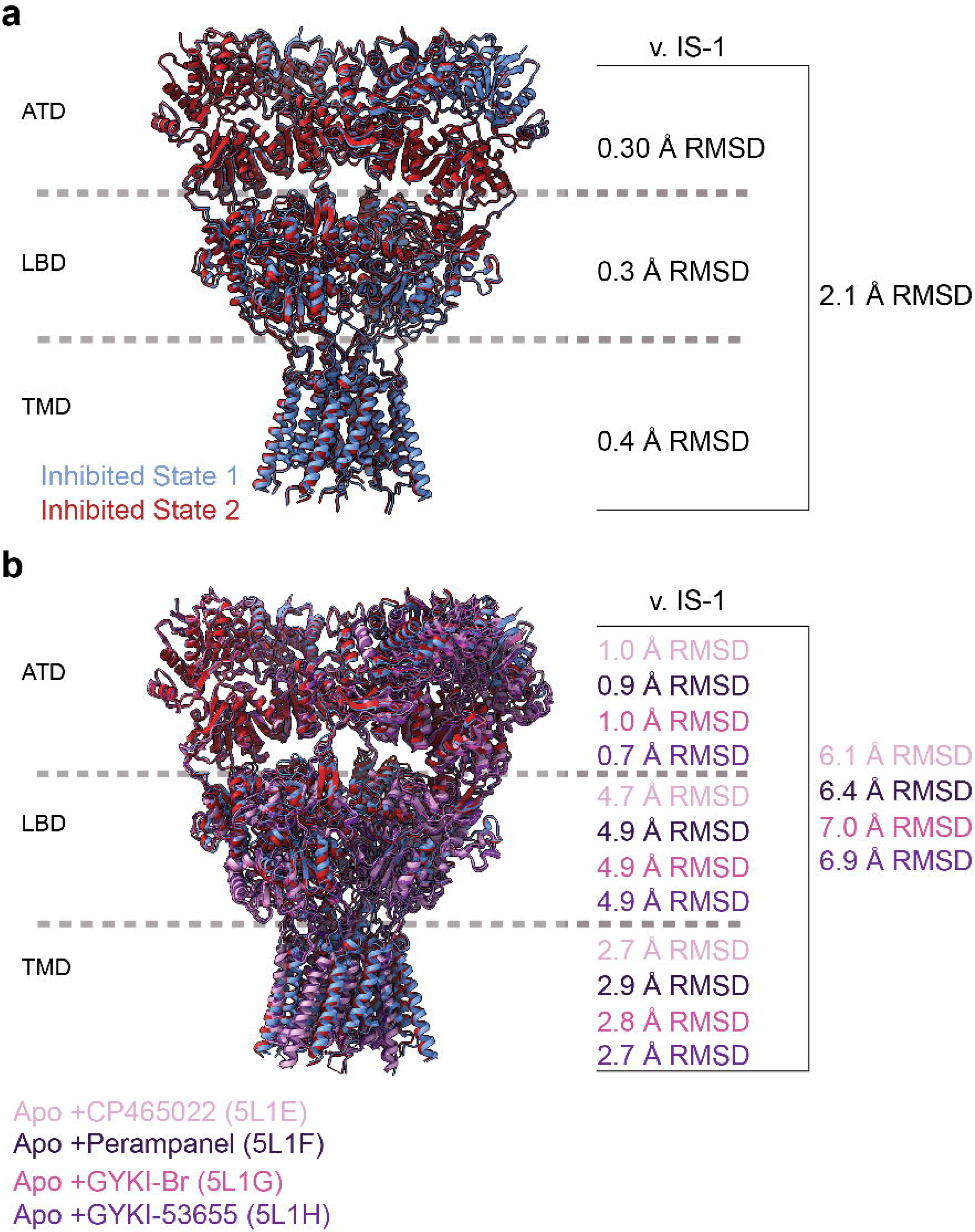
Comparison between inhibited states and AMPAR structures in resting state bound to PPLMs. a) overlay of IS-1 and IS-2 from this study demonstrating very minor deviation in the two states from each other. b) overlay of IS-1 and IS-2 against crystal structures of GluA2 in complex with PPLMs: CP465022 (PDB: 5L1E), Perampanel (PDB: 5L1F), GYKI-Br (PDB: 5L1G), and GYKI-53655 (PDB: 5L1H). RMSD is greatest within the LBD layer, in agreement with the conformational shifts observed following GluA2-TARPγ2 activation in the presence of GYKI-52466.

**Extended Data 3.**
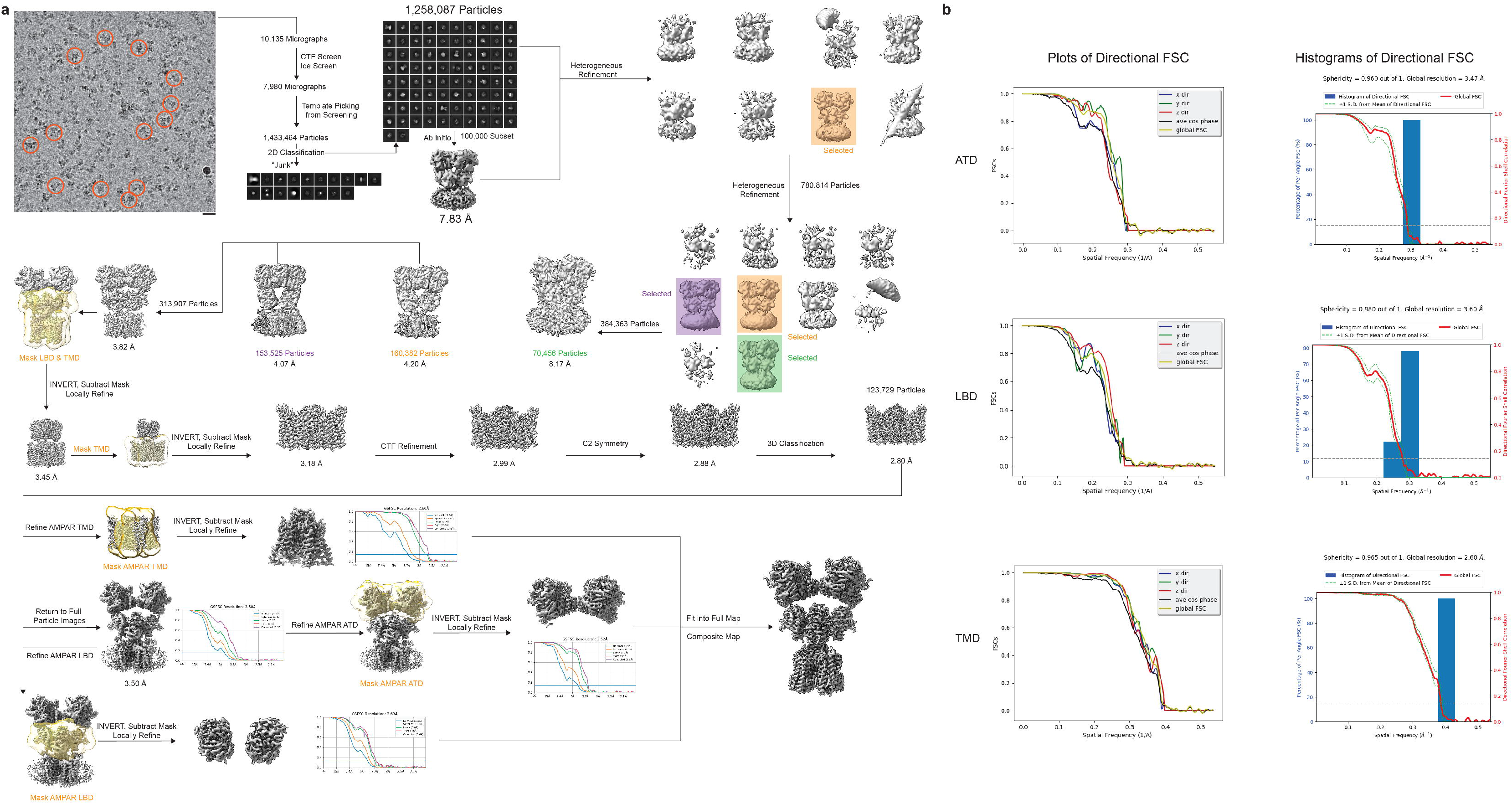
Cryo-EM processing workflow for IS-1.

**Extended Data 4.**
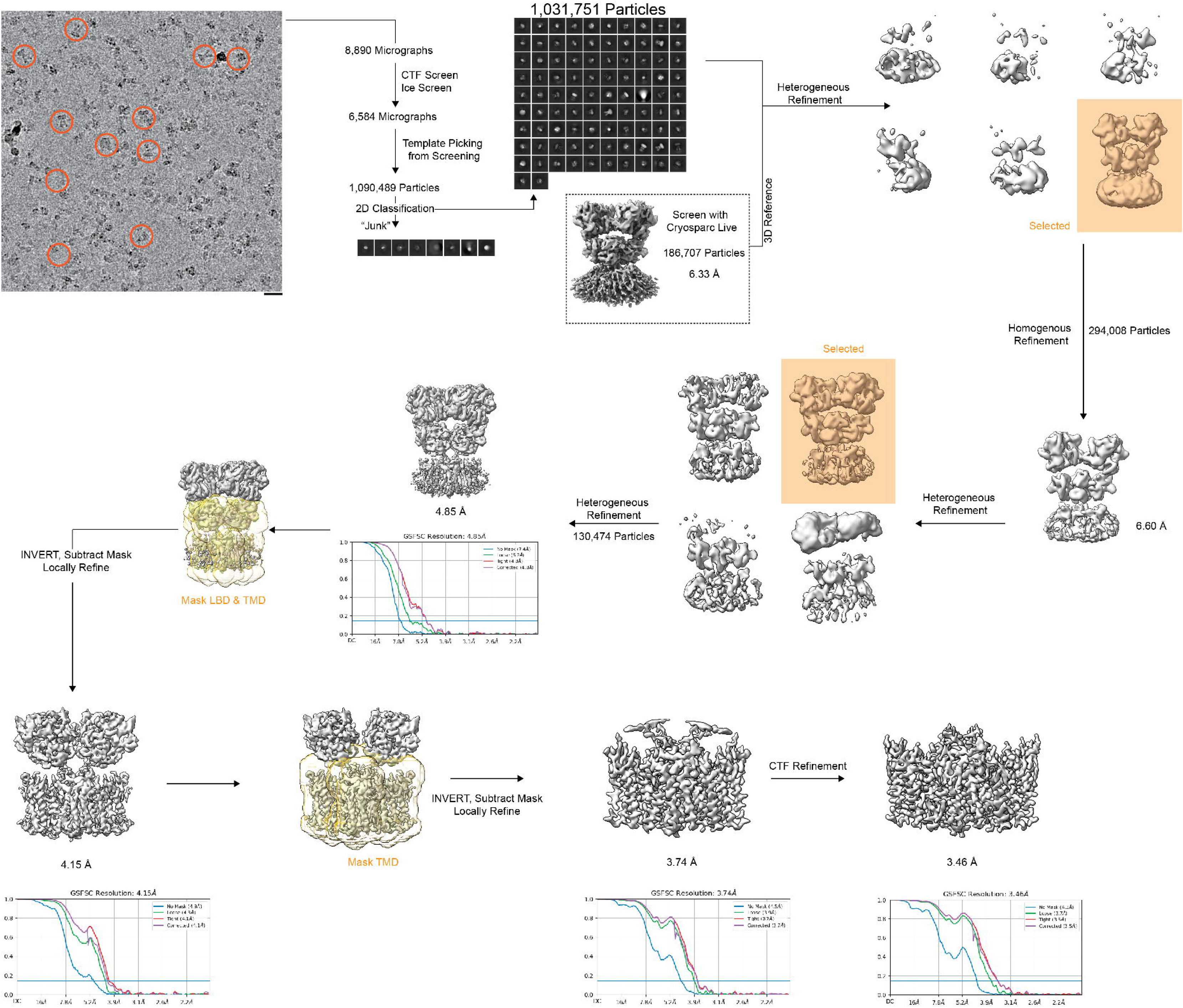
Cryo-EM processing workflow for IS-2.

**Extended Data 5.**
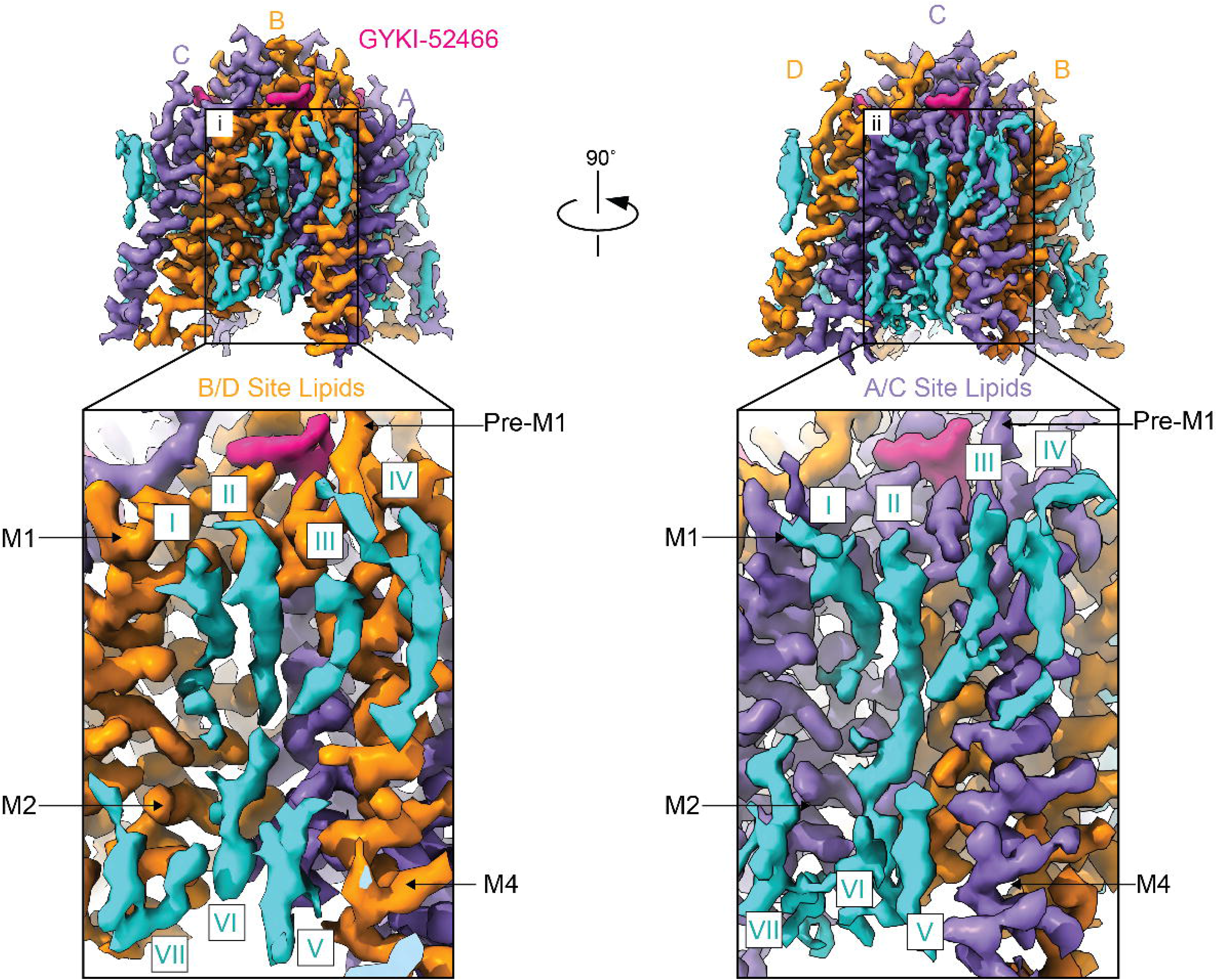
Structured lipids stabilize the AMPAR TMD. Coulomb potential maps of the AMPAR TMD from signal subtraction and focused refinement highlighting the presence of lipids (blue) bound to the TMD. (i) close-up of lipids bound to the B/D TMD subunits of the receptor showing at least seven distinct densities. (ii) close-up of the A/C subunits of the receptor demonstrating similar, but distinct lipid arrangements around the TMD.

**Extended Data 6.**
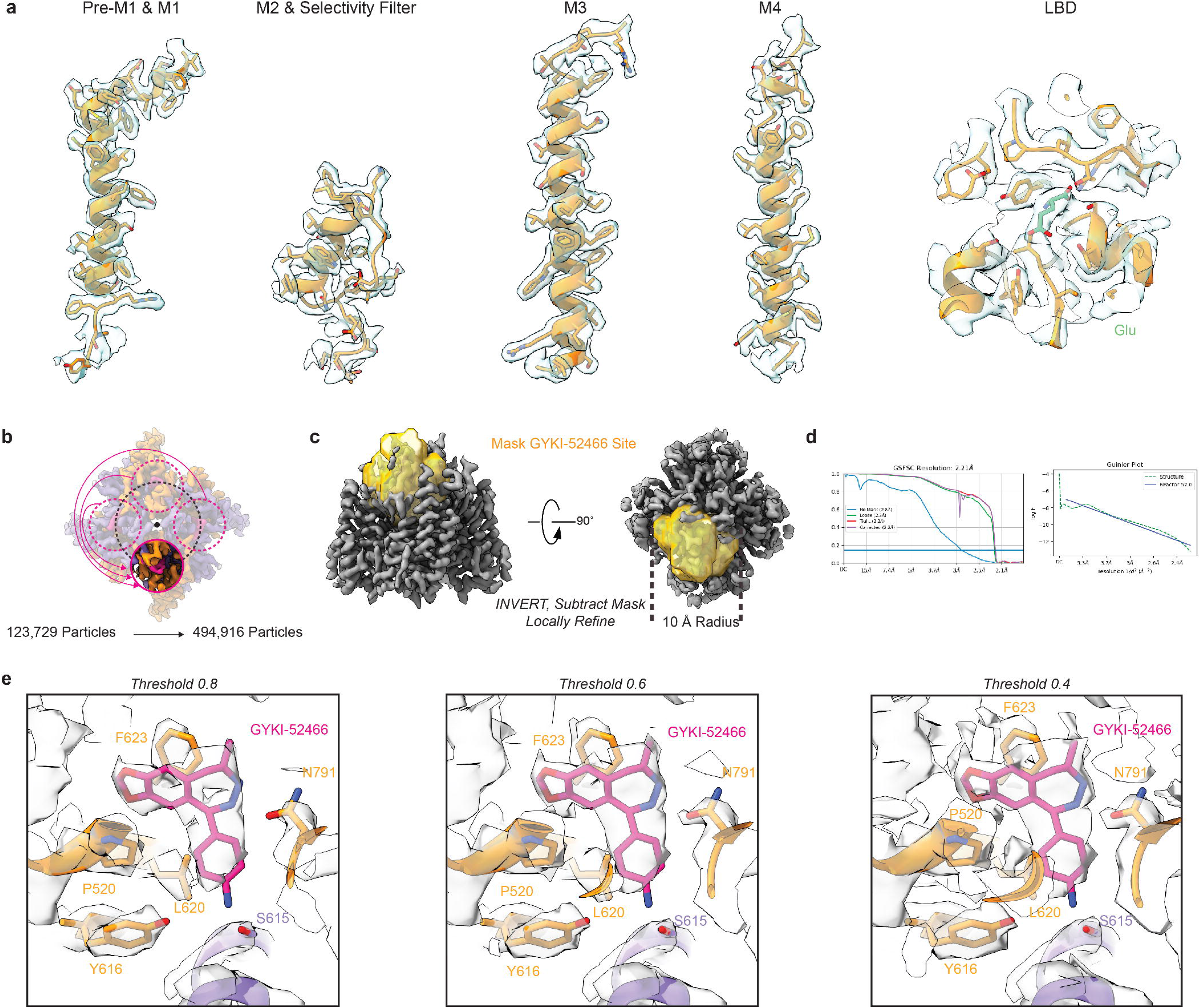
IS-1 Cryo-EM and workflow for elucidating GYKI-52466 binding pocket. a) Examples of Cryo-EM map for the AMPAR TMD and LBD. b) Symmetry expansion was applied through to the isolated IS-1 TMD to increase the effective particle count of the GYKI- 52466 binding pocket. b) Following expansion one of the four GYKI-52466 binding pockets was masked and then the mask inverted to subtract away the remaining TMD structure. c) Local refinement of the isolated GYKI-52466 binding pocket resolved the pocket to 2.21 Å resolution. d) Cryo-EM map of the GYKI-52466 shown from left to right at thresholds of 0.8, 0.6, and 0.4. Arrows point to potential water molecules within the binding pocket.

**Extended Data 7.**
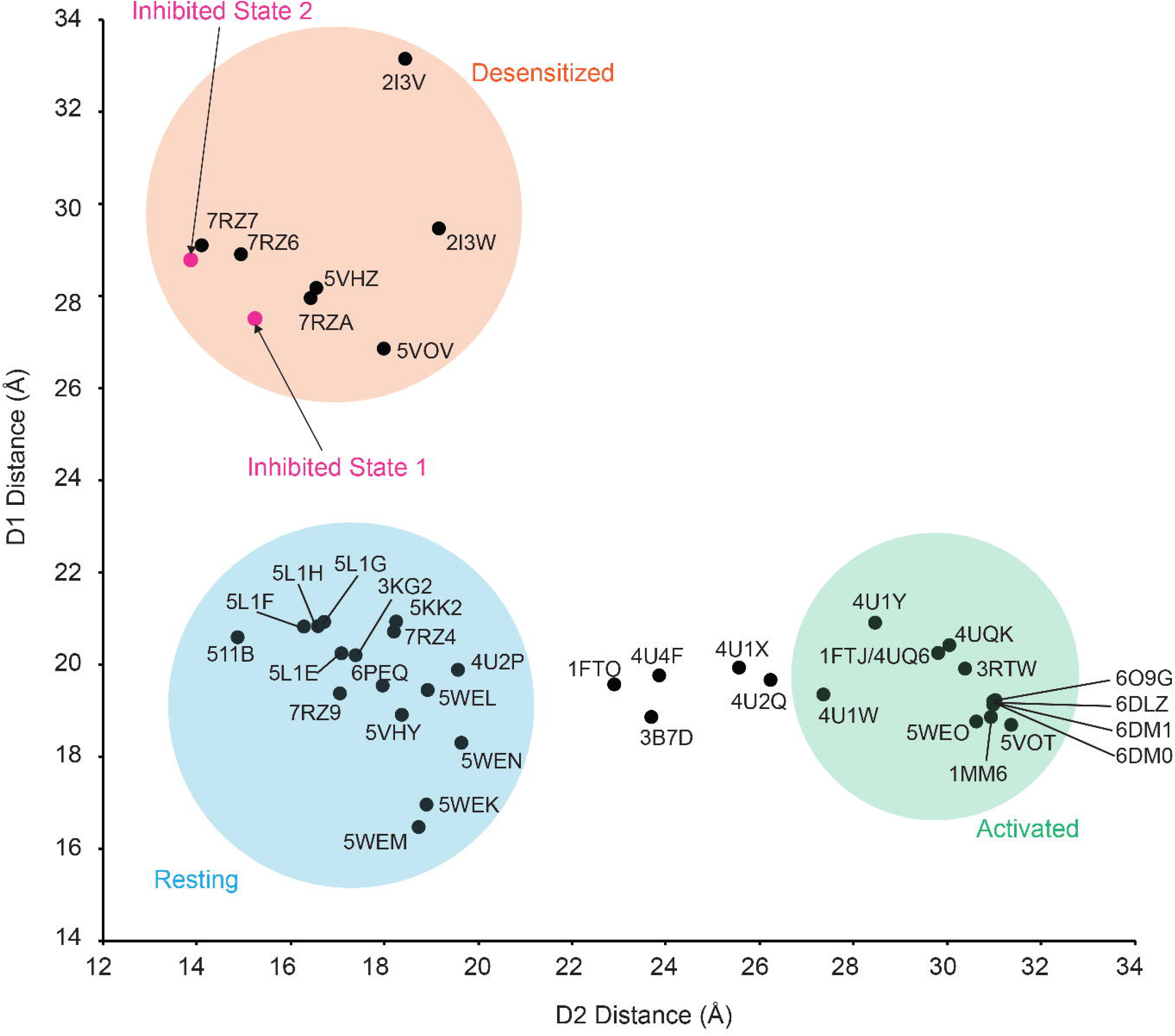
Clustering of AMPAR structures based on D1 and D2 distances. A detailed look of how published AMPAR structures cluster based on the measurements between the D1 and D2 lobes of LBD clamshells within local dimers. PDB reference numbers are given for each structure measured.

**Extended Data 8.**
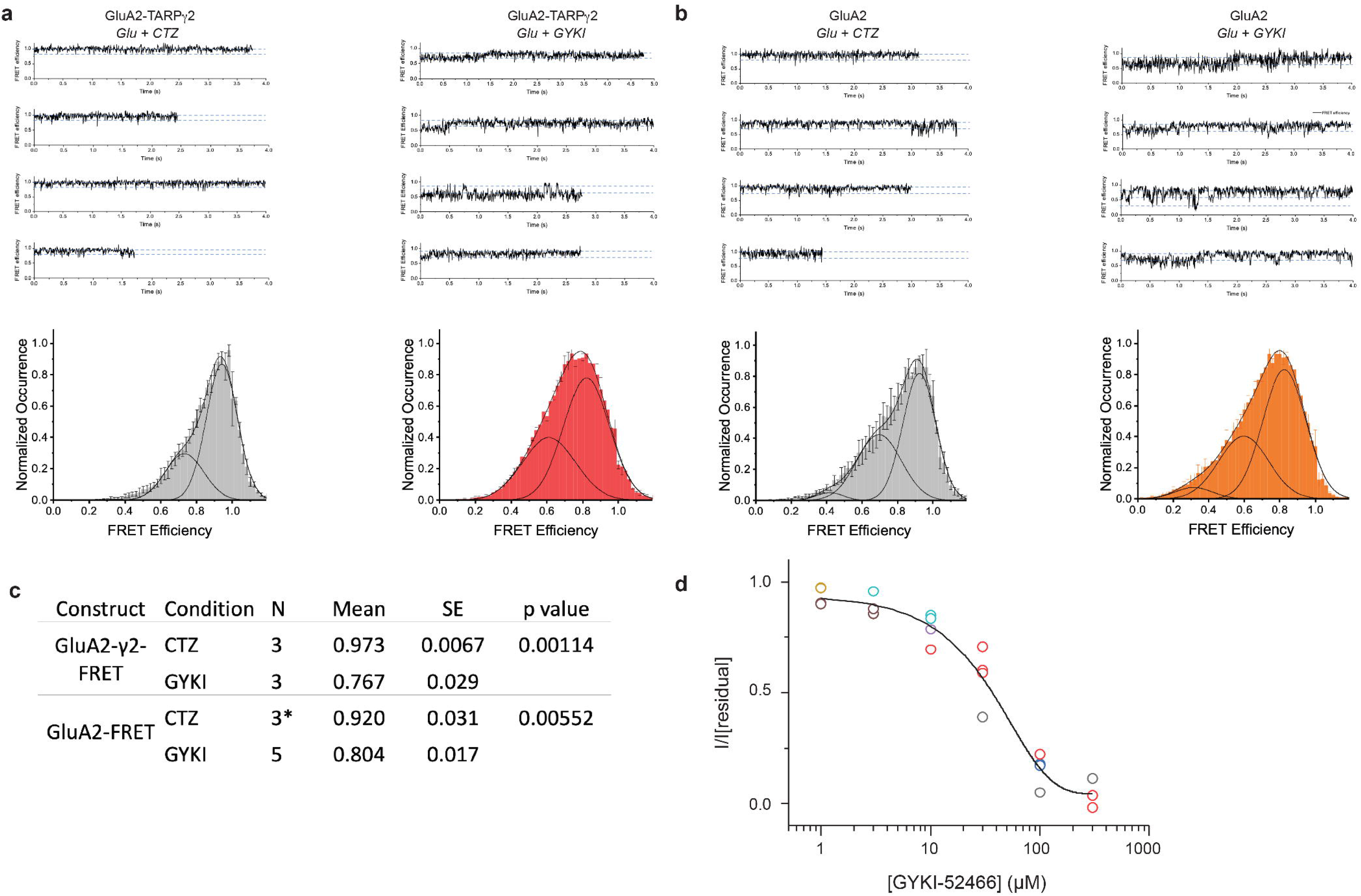
smFRET characterization of GYKI-52466 and CTZ allosteric modulation of AMPARs. a) (Top) Representative FRET efficiency traces for sampled four molecules in each condition (1 mM Glu + 100 µM CTZ or GYKI-52466) in GluA2-TARPγ2. (Bottom) FRET efficiency histogram generated from the compilation of all analyzed single-molecule traces. b) Same as panel a, but GluA2 without TARPγ2. c) Statistical analysis of the smFRET data. Mean of the mode for each day with standard error demonstrates a significant decrease in FRET efficiency from CTZ condition to GYKI-52466 condition using two-sample t-test assuming a one-tail distribution with known variances. *1 day of 30 molecules with 1mM of Glutamate and 100 μM CTZ were obtained from Carrillo and Shaikh et al. (2020). d) Concentration dependent inhibition by GYKI of GluA2-TARPγ2 residual currents in the presence of 1 mM of Glutamate and 100 μM of CTZ (IC50 = 39.87 ± 6.76 μM).

**Extended Data 9.**
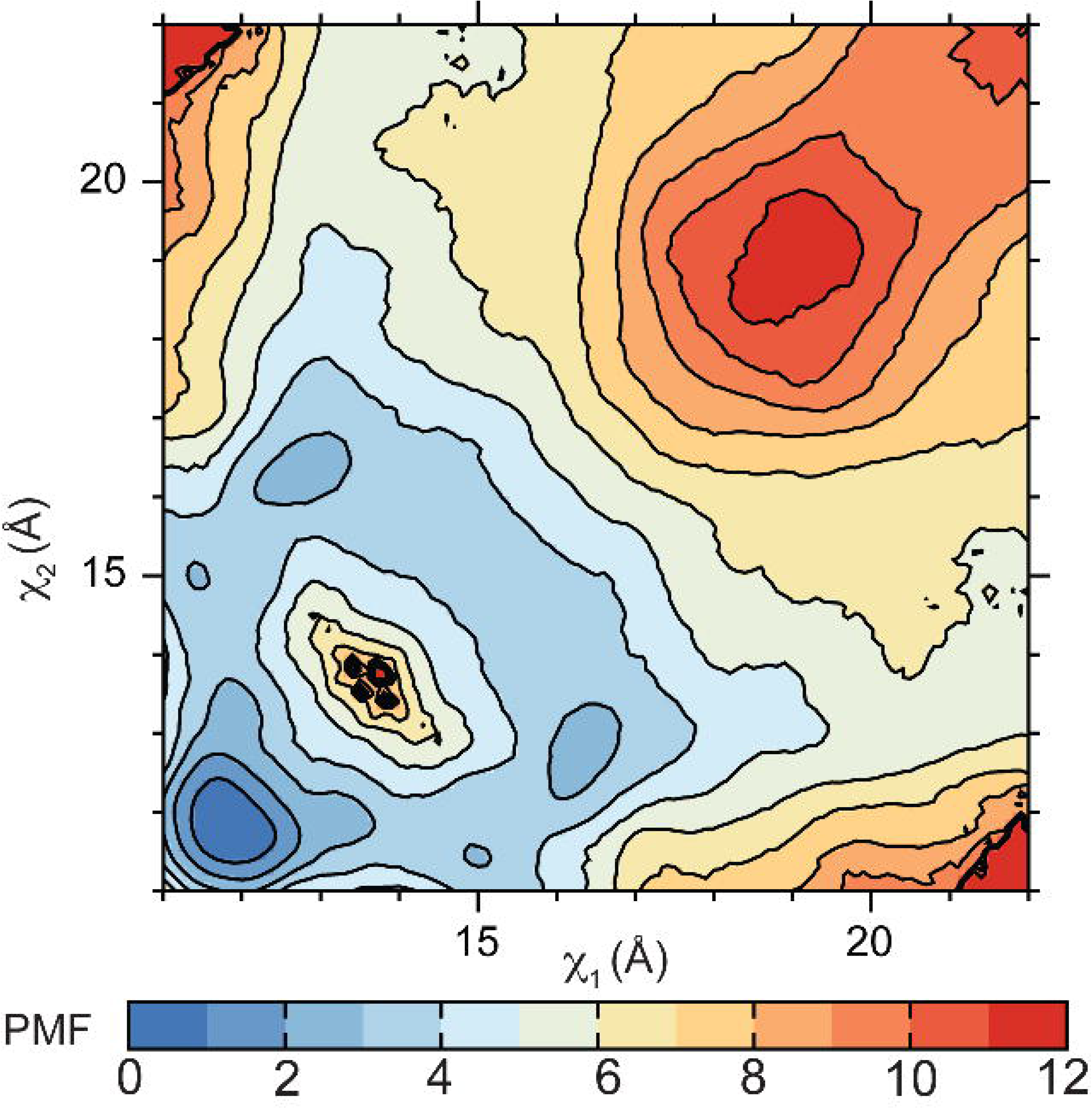
Free energy landscape governing desensitization in the GluA2-L483Y LBD dimer. The PMF is computed as a function of (χ_1_, χ_2_), the two distances between helices D and J at the dimer interface. The PMF is contoured in 1 kcal/mol increments.

**Extended Data 10.**
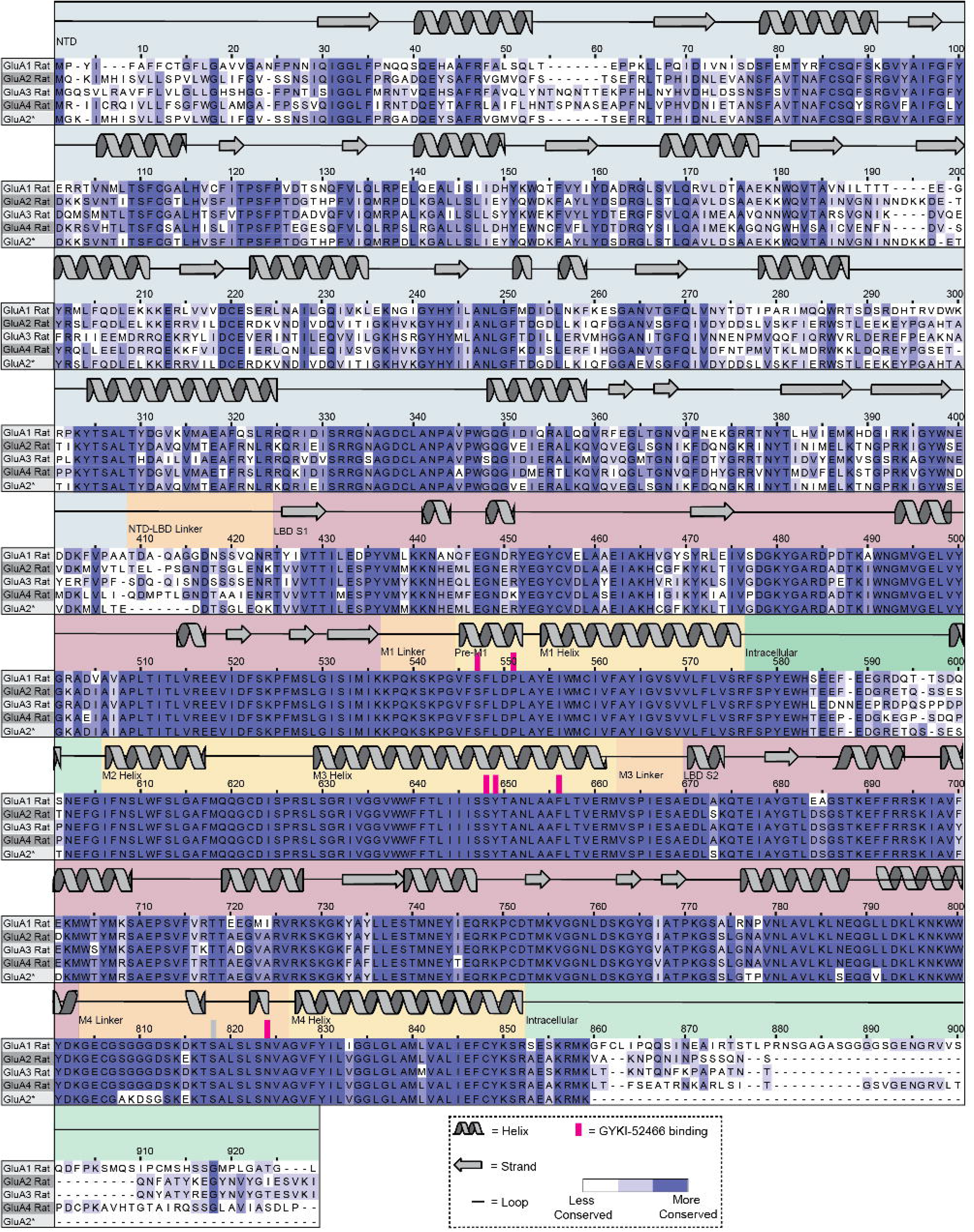
Alignment of GluA subunits. Multiple sequence alignment of Rat GluA1-4 protein sequences. Conservation is indicated by the intensity of purple coloring. Secondary structure is displayed above the alignment. GYKI-52466 interacting residues are highlighted in pink.

## Methods

### Construct design

The fusion construct GluA2-TARPγ2 was published previously^15,18,24,68^ and was constructed by fusing the GluA2 cDNA from rat to a GT linker, followed immediately by the N-terminus of mouse TARPγ2. The c-terminus of TARPγ2 is truncated following M4 followed by a Thrombin cleavage site, enhanced GFP (eGFP), and Strep Tag II immediately followed by a stop codon. This construct utilizes the pEG BacMam vector for baculovirus-driven protein expression in mammalian cells^69^.

For smFRET, the GluA2 construct was designed as previously described^33^. Briefly, cysteine residues 89, 196, and 436 were mutated to serines in the GluA2-flip (unedited Q isoform) construct to obtain the cysteine-light GluA2 construct, and a cysteine was introduced at position 467 for maleamide dye attachment to measure the intradimer interface of the LBD. To generate the GluA2-TARPγ2 construct for smFRET, the TARPγ8 from GluA2-TARPγ8 construct containing site L467C^33^ was replaced with TARPγ2 from the GluA2-TARPγ2 construct^36^ using restriction enzyme cloning with restriction enzymes BamHI and EcoRV to generate GluA2-TARPγ2 with cysteine site at 467.

### Protein expression and purification

GluA2-TARPγ2 bacmid was prepared as previously described^15,18,24^. P1 baculovirus was generated by transfecting ExpiSf9 cells (Gibco, A35243) cultured at 27 ˚C with polyethylenimine MW 40,000 (PolyScience, 24765). After 5 days, P1 virus was harvested, and expression in mammalian cells was induced by the addition of P1 baculovirus to Expi293F GnTI-cells (Gibco, A39240) grown in Expi293 media (Gibco, A14135101) in a 1:10 ratio of P1 virus to culture volume. Cells were grown in 37 ˚C, 5% CO_2_. 12-24 hours post-induction, the cell culture media was brought up to 10 mM Sodium Butyrate (Sigma, 303410) and 2 µM ZK 20075 (Tocris, 2345) and moved into a 30 ˚C, 5% CO_2_ incubator. The cells were harvested 72 h after transduction by centrifugation (5,000 *g*, 20 min at 4 ˚C) washed with PBS (pH 7.4) with protease inhibitors added (0.8 µM aprotinin, 2 µg ml^-1^ leupeptin, 2 µM pepstatin A, and 1 mM phenyl-methylsulfonyl fluoride) and then pelleted again (4800 *g*, 10 min at 4 ˚C). Supernatant was discarded and pellets were stored at -80 ˚C until purification. Pellets were thawed rotating in Lysis Buffer (150 mM NaCl, 20 mM Tris pH 8.0) with protease inhibitors added. Cells were lysed in an ice bath with a blunt probe sonicator (3 cycles, 1s on, 1s for 1 min, 20 W power). Lysed cells were centrifuged to pellet large cellular debris (4,800 *g*, 20 min at 4 ˚C). Supernatant was ultracentrifuged to pellet membranes (125,000 *g*, 45min), which were solubilized in solubilization buffer (150 mM NaCl, 20 mM Tris pH 8.0, 1% n-Dodecyl-β-D-Maltopyranoside (Anatrace, D310) and 0.2% Cholesteryl Hemisuccinate Tris salt (Anatrace, CH210) for 2 hours at 4°C under constant stirring. Insoluble material was pelleted in an ultracentrifuge (125,000 *g*, 45min at 4 ˚C) and solubilized protein was incubated with 0.75 mL Streptactin XT 4Flow resin (iba, 2-5010) per 1 L of cells overnight, rotating at 4°C. The following day, the resin was collected via gravity flow, and the resin was washed with 20 column volumes of GDN buffer (150 mM NaCl, 20 mM Tris pH 8.0, 0.01% GDN (glycol-diosgenin, Anatrace, GDN101)), before elution in GDN buffer made up to 50 mM D-biotin. Eluate was collected in a centrifugal concentrator and concentrated into a 500 µL volume at 4 ˚C. To remove eGFP and Strep Tag II, the concentrated protein was incubated with thrombin (1:200 w/w) for 1 h at 22 ˚C. The cleavage reaction was separated over a Superose 6 increase 10/300 column (Cytiva, 29091596) using an AKTA FPLC in GDN buffer. Peak fractions were collected and concentrated to 4.5 mg/ml.

### Sample preparation and data collection

UltrAuFoil 300 mesh R 1.2/1.3 grids (Electron Microscopy Services, Q350AR13A) were plasma treated in a Pelco Easiglow (25 mA, 120 s glow time, 10 s hold time, (Ted Pella, 91000)). Purified sample was split into two conditions. IS-1 sample was made up to 100 µM cyclothiazide (CTZ, Tocris, 07-131-0) and spun in an ultracentrifuge to pellet insoluble material prior to preparation of grids (75,000 *g*, 45 min), whereas IS-2 sample was made up to 100 µM CTZ and 100 µM GYKI- 52466 (Tocris, 1454) prior to centrifugation (75,000 *g*, 45 min). IS-1 samples were spiked with 100 µM GYKI-52466) and 1 mM Glu (pH 7.4) immediately before application to grids. IS-2 samples were only spiked with 1 mM Glu before application to grids. In both cases, 3 µL of sample was applied to glow discharged grids in an FEI Vitrobot Mark IV (Thermo Fisher Scientific, wait time 10 s, blot force 5, blot time 4 s) at 8 ˚C and 100% humidity and plunge-frozen in liquid ethane. Grids were imaged with a 300-kV Titan Krios 3i microscope equipped with fringe free imaging, Falcon 4i camera, and Selectris energy filter set to 10 eV slit width. Micrographs were collected with a dose rate of 8.15 e^-^ pixel^-1^ s^-1^ and a total dose of 40.00 e^-^ Å^-2^. We collected 8,800 micrographs of the GYKI-1 condition (0.93Å/pix) and 7,900 micrographs of the GYKI-2 condition (0.93Å/pix). Automated collection was achieved with EPU software from ThermoFisher.

### Image processing

Cryosparc^70^ was used for all aspects of image processing. Refer to extended data figures 3 and 4 for details. The reconstruction quality was tested for anisotropic contribution to the Fourier shell correlation (FSC) with 3DFSC^71^.

### Model building, refinement, and structural analysis

All molecular modeling, refinement, and analysis were performed with a combination of ChimeraX^72^, Isolde^73^, Coot^74^, and Phenix^75,76^ made accessible through the SBgrid consortium^77^. As a starting model, the activate state GluA2-TARPγ2^15,68^ (PDB 5weo) was used. Each domain (ATD, LBD, and TMD) was isolated and rigid body fit into the IS-1 full-length cryo-EM reconstruction using ChimeraX. The rigid body position of each protomer was refined by isolating them within domain and rigid body fitting. Following, each domain was joined into a single model. The exact positioning of each amino acid was fine-tuned based on the locally refined map of each domain using Coot. Then, Isolde was used to refine the model, and GYKI-52466 was placed in the map with Coot and merged into the model. Phenix was used to refine the final model. To model IS-2, the IS-1 model was rigid body fit into the IS-2 reconstruction and refined with Isolde and Phenix. Model quality was assessed with MolProbity^78^. Visualizations and domain measurements were performed in ChimeraX. Pore measurements were made with MOLE Online^79^.

### Labeling, acquisition, and analysis for Single-Molecule FRET

HEK-293 cells overexpressing GluA2 or GluA2-TARPγ2 receptors were labeled with 1:4 ratio of maleimide derivatives of Alexa 555 (donor) and Alexa 647 (acceptor) fluorophores (Invitrogen) in extracellular buffer (135 mM NaCl, 3 mM KCl, 2 mM CaCl2, 20 mM glucose, and 20 mM HEPES, pH 7.4) at room temperature for 30 minutes. Post-labeling, the cells were washed and solubilized for 1 h at 4 °C with buffer containing 1% lauryl maltose neopentyl glycol (Anatrace, Maumee, OH, USA), 2 mM cholesteryl hydrogen succinate (MP Biomedicals, Irvine, CA, USA), and ¼ protease inhibitor tablet (Pierce™) in phosphate-buffer saline. Solubilized cells were filtered from insoluble debris by ultracentrifugation at 100,000× g for 1 h at 4 °C using a TLA 100.3 rotor.

For the slide preparation, we followed established experimental methods as previously described^80–84^. The coverslips were initially cleaned by bath sonication in Liquinox phosphate-free detergent (Fisher Scientific) and acetone treatment. Further cleaning involved incubating the slides in a 4.3% NH4OH and 4.3% H2O2 solution at 70 °C, followed by plasma cleaning using a Harrick Plasma PDC-32G Plasma Cleaner. The cleaned glass was aminosilanated using Vectabond reagent (Vector Laboratories), followed by polyethylene-glycol (PEG) treatment with 0.25% w/w 5 kDa biotin-terminated PEG (NOF Corp., Tokyo, Japan) and 25% w/w 5 kDa mPEG succinimidyl carbonate (Laysan Bio Inc., Arab, AL), then followed by a secondary PEG treatment with 25 mM short-chain 333 Da MS(PEG)4 Methyl-PEG-NHS-Ester Reagent (Thermo Scientific). A microfluidics chamber was constructed on the slide, comprising an input port, a sample chamber, and an output port. To coat the biotinylated surface with streptavidin molecules, 0.2 mg/mL Streptavidin in 1X smFRET imaging buffer (1 mM DDM (n-dodecyl-β-d-maltoside), 0.2 mM CHS (cholesteryl hydrogen succinate), and 1X phosphate-buffer saline was introduced into the chamber and incubated for 10 min before washing with 1X phosphate-buffer saline. 60μl of biotinylated Goat Anti-Mouse IgG (H + L) secondary antibody at 2.7ng/µl (Jackson Immunoresearch Laboratories, Inc., West Grove, PA, USA, catalog number 115-065-003) in 1X phosphate-buffer saline was then flowed through the chamber and incubated for 20 min, followed by 1x phosphate-buffer saline wash.

Following this, either 60μl of anti-GluR2 at 3ng/μl for GluA2-FRET purification, Clone: L21/32(BioLegend®) or 60μl of anti-TARPγ2 at 2.4ng/μl for GluA2-γ2-FRET purification, Clone: N245/36 (Millipore) in 1x phosphate-buffer saline was applied twice through the chamber and incubated for 20 min, followed by washing with 1x phosphate-buffer saline. Bovine serum albumin (0.1 mg/mL) was introduced into the chamber and incubated for 15 min, followed by 1x phosphate-buffer saline wash. Detergent-solubilized purified proteins were attached to the glass slide using an in situ immuno-precipitation method by applying 50 µL of sample three times through the chamber and incubating for 20 min. 90 µL of oxygen-scavenging solution buffer system (ROXS) was applied inside the chamber containing 1 mM methyl viologen, 1 mM ascorbic acid, 0.01% w/w pyranose oxidase, 0.001% w/v catalase, 3.3% w/w glucose (all from Sigma-Aldrich, Inc., St. Louis, MO, USA), 1 mM DDM (Chem-Impex, Wood Dale, IL, USA), and 0.2 mM CHS (MP Biomedicals, LLC, Santa Ana, CA, USA) in phosphate-buffer saline, pH 7.4. For CTZ condition, 1mM glutamate, and 100 μM of CTZ were introduced into the ROXS. In the GYKI-52466 treated condition, 1mM glutamate and 100 µM GYKI-52466 (MilliporeSigma, Burlington, MA, USA), were introduced into the ROXS.

The smFRET data was collected using a MicroTime 200 Fluorescence Lifetime Microscope from PicoQuant. A donor excitation laser (532 nm; LDH-D-TA-530; Picoquant, Berlin, Germany) and an acceptor excitation laser (637 nm; LDH-D-C-640; Picoquant) were employed, utilizing a Pulsed Interleaved Excitation (PIE) scheme to excite the fluorophores. Emitted photons were collected through the objective lens (100X 1.4 numerical aperture; Olympus). Emission filters for the donor (550 nm; FF01-582/64; AHF, Tübingen-Pfrondorf, Germany or Semrock, Rochester, NY) and acceptor (650 nm 2XH690/70; AHF) were used to select photons for each detection channel. These photons were directed to two SPAD photodiodes (SPCM CD3516H, Excelitas technologies, Waltham, MA) to measure the fluorescence intensity for each fluorophore. The donor and acceptor fluorescence intensities were recorded for one protein at a time.

In our data analysis, we selected only those molecules that exhibited a single photobleaching step in both the donor and acceptor channels. This stringent criterion ensured that only one donor and one acceptor fluorophore were attached to each GluA2 protein. Furthermore, we retained only those molecules that displayed anti-correlation between the donor and acceptor fluorescence, confirming that the fluorophores were engaged in FRET prior to photobleaching. Molecules not exhibiting these characteristics were excluded from the final analysis. The number of molecules included in the analysis for each condition is as follows: GluA2-γ2-FRET (CTZ = 76, GYKI-52466 = 77), GluA2-FRET (CTZ = 62*, GYKI-52466 = 96). *30 molecules with 1mM of glutamate and 100μM CTZ were obtained from Carrillo and Shaikh et al. (2020)^33^.

The corrected donor and acceptor intensities over time were then used to calculate a FRET efficiency trace for each molecule. These traces were pooled for each condition and used to create FRET efficiency distribution histograms for each condition. We conducted Step Transition and State Identification (STaSI) analysis to determine the number of conformational states in each condition^85^. The smallest number of states that accurately described the data as determined by the STaSI analysis was adopted as the final number of states for each condition. Using the results of the STaSI analysis and Origin software (OriginLab), the FRET efficiency histograms for each condition were fitted with Gaussian curves to represent the conformational states within the overall distributions.

To test for the statistical difference between conditions CTZ and GYKI, FRET efficiency mode was obtained for each day as this more accurately represents the histogram peak. The mean and standard deviation were calculated across these days. A two-sample t-test, assuming a one-tail distribution with known variances, was used to assess the statistical differences between the conditions using Origin software (OriginLab).

### Electrophysiology

For electrophysiological measurements of GluA2-TARPγ2, 1μg of DNA was transfected in 3cm culture dishes using Lipofectamine 2000. Patch-clamp recordings were performed 24–48 h after transfection using fire-polished borosilicate glass (Sutter instruments, Novato, CA, USA) pipettes with 1–4 megaohms resistance were filled with internal solution: 110 mM CsF, 30 mM CsCl, 4 mM NaCl, 0.5 mM CaCl2, 10 mM HEPES, and 5 mM EGTA (adjusted to pH 7.4 with CsOH). The extracellular solution consisted of 150 mM NaCl 3 mM KCl, 2 mM CaCl2, and 10 mM HEPES adjusted to pH 7.4 with NaOH. External solutions were locally applied to lifted cells or patches using a SF-77B perfusion fast-step (Warner Instruments, Holliston, MA, USA). For inhibition-dose response, 1 mM of glutamate and 100 μM CTZ was applied first to obtain control condition followed by 2-3 different concentrations of GYKI with 1 mM of glutamate. For each GYKI concentration, the same GYKI concentration was preincubated in extracellular buffer. Recordings were performed using an Axopatch 200B amplifier (Molecular Devices, San Jose, CA, USA) at −60 mV hold potential, acquired at 2kHz using pCLAMP10 software (Molecular Devices, Axon 200B and Digidata 1550A; Molecular Devices). Individual patch-clamp traces were analyzed, and data was reduced to 300μs sampling using Clampfit 11 software (Molecular Devices, San Jose, CA, USA). IC50 was quantified from the average residual current using Clampfit 11. Representative traces graphed, normalized, and calculated using Origin software (OriginLab).

### Free Energy Molecular Dynamics Simulations

The conformational free energy landscape, or potential of mean force (PMF), of the LBD dimer was computed using umbrella sampling simulations. A two-dimensional order parameter (χ_1_, χ_2_) describes large-scale conformational transitions between each LBD of the dimer. χ_1_ and χ_2_ each indicates the distance between the center of mass (COM) of the atoms N, CA, CB, C, and O in residues 482–488, helix D, and the COM of the same atoms in residues 748–757, helix J. Helices D and J form the dimer interface. Coordinates for the umbrella sampling windows were generated via targeted (biased-potential) molecular dynamics (MD) simulations using CHARMM^86^ in 1 Å increments along χ_1_ and χ_2_. These coordinates were initiated from the crystal structure of a glutamate-bound GluA2 LBD dimer (PDB ID 1FTJ)^45^. For GluA2-L483Y, these coordinates were initiated from the crystal structure of the mutant LBD dimer (PDB: 1LB8)^22^. Missing residues were built using the ModLoop server^87^, and missing residue side chains were built using SCWRL4^88^.

All simulations were performed using CHARMM36 with explicit solvent at 300 K. The all-atom potential-energy function PARAM27 for proteins^89,90^ and the TIP3P potential-energy function for water^91^ were used. Each simulation system contains ∼56,000 atoms, and 39 Na^+^ and 47 Cl^−^ ions were added in the bulk solution to give ∼150 mM NaCl and an electrically neutral system. Periodic boundary conditions were used with an orthorhombic cell with approximate dimensions 96 Å x 78 Å x 78 Å. Equilibration was carried out in the NVT ensemble with restraints applied to the backbone and sidechain atoms, which were slowly released over the course of the equilibration. Production simulations were carried out in the NPT ensemble at 1 atm and 300 K^92^. Long-range electrostatic interactions were computed using the particle mesh Ewald (PME) algorithm^93^.

The PMF comprises 140 umbrella sampling windows totaling 364 ns of simulation time and 398 ns for GluA2-L483Y. Harmonic biasing potentials with a force constant of 2 kcal mol^−1^ Å^−2^ centered on (χ_1_, χ_2_) were used. Each PMF was computed using the weighted histogram analysis method (WHAM)^94,95^ to unbias and recombine the sampled distribution functions from all windows.

## Data Availability

All cryo-EM reconstructions will be deposited into the Electron Microscopy Data Bank (EMDB) upon publication. All micrographs from the IS-1 and IS-2 datasets will be deposited into the Electron Microscopy Public Image Archive (EMPIAR) upon publication. All structural models generated from cryo-EM will be deposited in the Protein Data Bank upon publication. All conformers from MD simulation trajectories, data from umbrella sampling, analysis code, will be publicly available from Zenodo upon publication of this work.

## Acknowledgements

We thank members of the Twomey, Huganir, Lau, and Jayaraman labs for insightful discussions and L. Dillard (Twomey lab) for assistance in collecting the IS-2 dataset. We thank M. Catipovic (JHU) for insightful comments on the manuscript. All cryo-EM data was collected at the Beckman Center for Cryo-EM at Johns Hopkins with assistance from D. Sousa and D. Ding. Computational resources were provided by the Maryland Advanced Research Computing Center (MARCC) and Advanced Research Computing at Hopkins (ARCH) at Johns Hopkins University.

## Author Contributions

E.C.T. and R.L.H. supervised all aspects and planning of this research. E.C.T. and W.D.H. designed the project. E.C.T. and W.D.H. wrote the manuscript with input from all authors. W.D.H. prepared samples for cryo-EM, collected cryo-EM data, processed cryo-EM data, analyzed data, and built models with E.C.T. A.M.R. assisted with protein expression, model building, data analysis, structural analysis, and in uncovering the inhibition mechanism with W.D.H. V.J., C.U.G. designed the smFRET and electrophysiology experiments with input from E.C.T. and W.D.H. C.U.G. carried out the smFRET and electrophysiology experiments under the supervision of V.J. C.U.G. carried out statistical analysis of smFRET and electrophysiology data under supervision of V.J. A.Y.L. planned and carried out all molecular dynamics simulation studies and analysis.

Cryo-EM data collection, refinement, and validation statistics.

1 Cryo-EM data collection, refinement, and validation statistics
|  | IS-1<br>(EMDB-xxxx)<br>(PDB xxxx) | IS-2<br>(EMDB-xxxx)<br>(PDB xxxx) |
| --- | --- | --- |
| **Data collection and processing** |  |  |
| Magnification | 130,000x | 130,000x |
| Voltage (kV) | 300 | 300 |
| Electron exposure (e-/Å<sup>2</sup>) | 40 | 40 |
| Defocus range (μm) | -1.0 – 2.6 | -1.0 – 2.6 |
| Pixel size (Å) | 0.93 | 0.93 |
| Symmetry imposed | C2 | C2 |
| Initial particle images (no.) | 1,258,087 | 1,031,751 |
| Final particle images (no.) | 123,729 | 130,474 |
| Map resolution (Å) |  |  |
| FSC = 0.143 |  |  |
| Map resolution range (Å) | 2 – 13 | 2.5 – 13 |
| **Refinement** |  |  |
| Initial model used (PDB code) | 5WEO | This study, IS-1 |
| Model resolution (Å) | 4.2 | 3.55 |
| FSC = 0.143 |  |  |
| Model resolution range (Å) | 3.4 – 4.1 | 3.2 – 4.1 |
| Map sharpening *B* factor (Å<sup>2</sup>) | -65 | -120 |
| Model composition |  |  |
| Non-hydrogen atoms | 25180 | 25180 |
| Protein residues | 3186 | 3186 |
| Ligands | 4 | 4 |
| *B* factors (Å<sup>2</sup>) |  |  |
| Protein | 0.48/164.05/76.91 | 0/391.17/121.95 |
| Ligand | 12.19/43.65/29.89 | 0.01/9.83/4.8 |
| R.m.s. deviations |  |  |
| Bond lengths (Å) | 0.003 | 0.004 |
| Bond angles (°) | 0.641 | 0.651 |
| Validation |  |  |
| MolProbity score | 1.7 | 1.66 |
| Clashscore | 6.58 | 7.04 |
| Poor rotamers (%) | 1.33 | 0.34 |
| Ramachandran plot |  |  |
| Favored (%) | 96.3 | 96.05 |
| Allowed (%) | 3.51 | 3.77 |
| Disallowed (%) | 0.19 | 0.18 |

